# A mechanism of synergistic Mediator recruitment in RNA polymerase II transcription activation revealed by single-molecule fluorescence

**DOI:** 10.1101/2024.12.10.627625

**Authors:** Daniel H. Zhou, Jongcheol Jeon, Nida Farheen, Larry J. Friedman, Jane Kondev, Stephen Buratowski, Jeff Gelles

## Abstract

Transcription activators trigger transcript production by RNA Polymerase II (RNApII) via the Mediator coactivator complex. Here the dynamics of activator, Mediator, and RNApII binding at promoter DNA were analyzed using multi-wavelength single-molecule microscopy of fluorescently labeled proteins in budding yeast nuclear extract. Binding of Mediator and RNApII to the template required activator and an upstream activator sequence (UAS), but not a core promoter. While Mediator and RNApII sometimes bind as a pre-formed complex, more commonly Mediator binds first and subsequently recruits RNApII to form a preinitiation complex precursor (pre-PIC) tethered to activators on the UAS. Interestingly, Mediator occupancy has a highly non-linear response to activator concentration, and fluorescence intensity measurements show Mediator preferentially associates with templates having at least two activators bound. Statistical mechanical modeling suggests this “synergy” is not due to cooperative binding between activators, but instead occurs when multiple DNA-bound activator molecules simultaneously interact with a single Mediator.

## Introduction

Eukaryotic genes respond to a wide range of cues. Multicellular organisms contain many differentiated cell types, each expressing distinct subsets of the same genomic information^1,2^. Even single-cell eukaryotes such as yeast exhibit different transcription patterns tuned to nutrient availability, temperature, and other environmental conditions. Mediating these complex patterns of gene expression are transcription activators (TAs), sequence-specific DNA binding proteins that stimulate the RNA Polymerase II (RNApII) transcription machinery to assemble pre-initiation complexes (PICs) on target promoters^3–5^. A more complete understanding of transcription activation mechanisms remains an essential goal.

Binding sites for TAs are typically clustered within DNA elements known as enhancers or upstream activating sequences (UASs)^6^. Metazoan enhancers often have a dozen or more TA binding sites^1^. Importantly, two or more binding sites for the same TA can produce far greater activation than the sum of the individual sites, a phenomenon termed “synergy”^7–9^. Moreover, enhancers with sites for different TAs can synergistically integrate their different signaling responses, providing a basis for combinatorial regulation ^4,10–12^. However, multiple TA sites in an enhancer do not always produce synergy^12–15^, and multiple mechanisms for synergy have been proposed^9,14,16–19^.

TAs function through multiple coactivators. Chromatin modifying and remodeling coactivators (e.g., SAGA, SWI/SNF) modulate promoter accessibility. However, the most critical coactivator may be Mediator^20–24^. This large, multi-subunit complex ^24–30^ is proposed to “mediate” communication between UAS-bound TAs and PIC components at the core promoter, consistent with chromatin immunoprecipitation experiments detecting Mediator at either enhancers or core promoters depending on circumstances^28,31–34^. Likewise, structural studies show Mediator either incorporated into the PIC at the core promoter^35–39^ or bound with TAs, RNApII, and some general transcription factors (GTFs) at the UAS^40^. Finally, imaging in vivo led to models where multiple Mediator molecules form condensates at enhancers via intrinsically disordered regions^41,42^. Given this diversity of interactions, structural poses, and functions, the molecular mechanisms by which Mediator conveys activating signals from TAs to the PIC machinery at core promoters are still not fully understood^31,43–45^.

Systems comprising multiple macromolecular interactions are often best studied by single-molecule methods. We used colocalization single-molecule spectroscopy (CoSMoS)^46,47^ to directly image TA, RNApII, and Mediator dynamics on individual DNA molecules in real time. Our *Saccharomyces cerevisiae* nuclear extract system contains the full complement of nuclear proteins and recapitulates essential features of activated transcription in vitro^48–51^ when supplemented with the recombinant TA Gal4-VP16^52–54^. To focus solely on chromatin-independent transcription activation mechanisms, the template DNA was not pre-assembled into chromatin.

This system previously revealed that RNApII and GTFs do not sequentially assemble PICs directly on the core promoter as often assumed^49^. Instead, RNApII, TFIIF, and TFIIE associate first at the TA-bound UAS to form a PIC precursor (pre-PIC), which we proposed then transfers to the core promoter. Here we show that Mediator also initially binds UAS-associated TAs independently of the core promoter, and that this binding is necessary for simultaneous or subsequent RNApII binding. Quantitative analysis of Mediator and TA occupancy on DNA reveals that a single Mediator strongly prefers templates bound by multiple TAs, implicating Mediator recruitment as a locus of TA synergy.

## Results

### Dynamics of Gal4-VP16 and Mediator binding to DNA

To characterize how UAS-bound TAs affect Mediator dynamics, TA binding was monitored using recombinant Gal4-SNAP-VP16 protein consisting of the yeast Gal4 DNA-binding domain, a SNAP tag fluorescently labeled with red-excited dye adduct SNAP-Surface-649, and the potent herpes simplex virus VP16 transactivation domain (TAD)^51,52^ (**Figure S1A**). This protein is hereafter designated Gal4-VP16^649^. The DNA template contains a UAS comprised of five Gal4 binding sites positioned upstream of the *CYC1* core promoter (UAS+promoter; **Figure 1A**; **Table S4**). Transcription from this template in yeast nuclear extract (YNE) is as strongly activated by Gal4-SNAP-VP16^51^ as by Gal4-VP16^49,50,55,56^. Extracts were prepared from yeast grown in glucose medium and thus lack endogenous Gal4 activity^57,58^.

**Figure 1:**
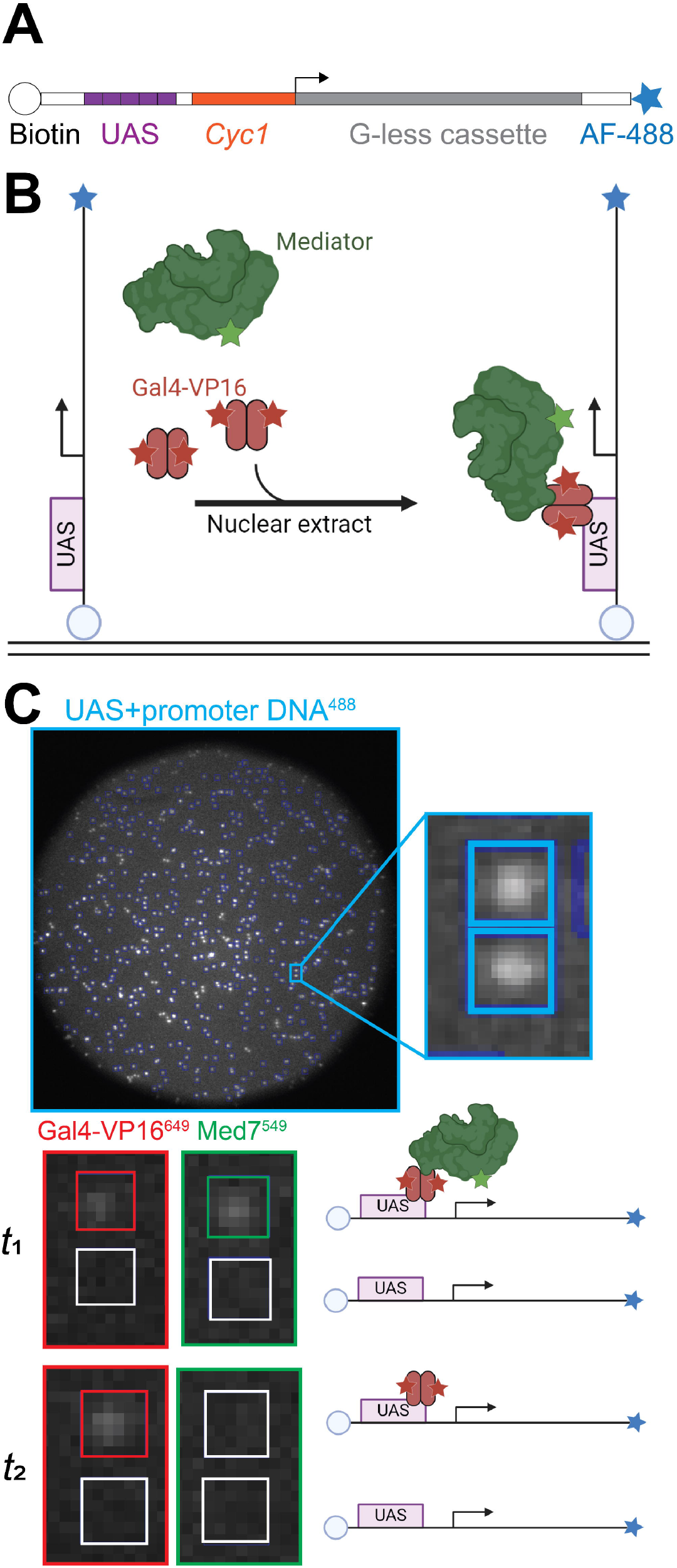
CoSMoS experiments design and example data. **(A)** Schematic of the UAS+promoter DNA. **(B)** Experimental design (not to scale). Labeled Gal4-VP16^649^ (red) and Med7^549^ (green) in YNE were allowed to bind to dye-labeled UAS+promoter DNA tethered to the slide surface. **(C)** Single-molecule imaging example. Large image shows the microscope field of view (65 µm diameter) before addition of extract at time *t* = 0 with locations of DNA molecules marked (blue). Insets show a magnified region with two DNA molecules (boxes). Different laser excitation wavelengths visualized the DNA (blue border; *t* < 0) and colocalized Gal4-VP16^649^ (red border) or Med7^549^ (green border) at times *t*_1_ = 101 s and *t*_2_ = 301 s after extract addition. Cartoons illustrate inferred molecular species present on the two DNA molecules at *t*_1_ and *t*_2_. See also **Figure S1** and **Table S4**.

To observe Mediator, *S. cerevisiae* strain YSB3613 (**Table S1**) was constructed with tandem HA and SNAP tags fused to the Med7 C-terminus. Med7 is essential for viability^38^, so the unperturbed growth of this strain (**Figure S1B**) suggests that the tagged Mediator functions normally *in vivo*. Immunoblotting confirmed expression of Med7-HA-SNAP protein at the expected molecular weight (**Figure S1C**), and YNE treated with the green-excited dye adduct SNAP-Surface-549 shows a single fluorescent protein (hereafter designated Med7^549^) at the same position **(Figure S1D**). Bulk *in vitro* transcription assays confirmed that Med7^549^-containing YNE exhibits activator-dependent transcription (**Figure S1E**).

Single-molecule microscopy experiments were done as previously described^49–51^ (**Figure 1B**). UAS+promoter DNA templates, tagged with biotin and blue-excited dye AF488, were tethered in a microscope flow chamber^46,47^. Total internal reflection fluorescence (TIRF) microscopy revealed several hundred DNAs in a typical field of view (**Figure 1C**). These were incubated with Med7^549^-containing YNE and Gal4-VP16^649^ in the absence of NTPs, conditions allowing stable formation of transcription-competent PICs while preventing transcription initiation^48–50,55^.

At 10 nM Gal4-VP16^649^, binding was seen at 74% (376 of 506) of DNA locations (**Figure 2A, S2A** upper panel). TA molecules remained bound for up to hundreds of seconds. In contrast, only 8% (28 of 356) of randomly selected control locations lacking labeled DNA molecules displayed TA binding (**Figure 2B** and **Figure S2A** lower panel). Thus, TA bound specifically to template DNA.

**Figure 2:**
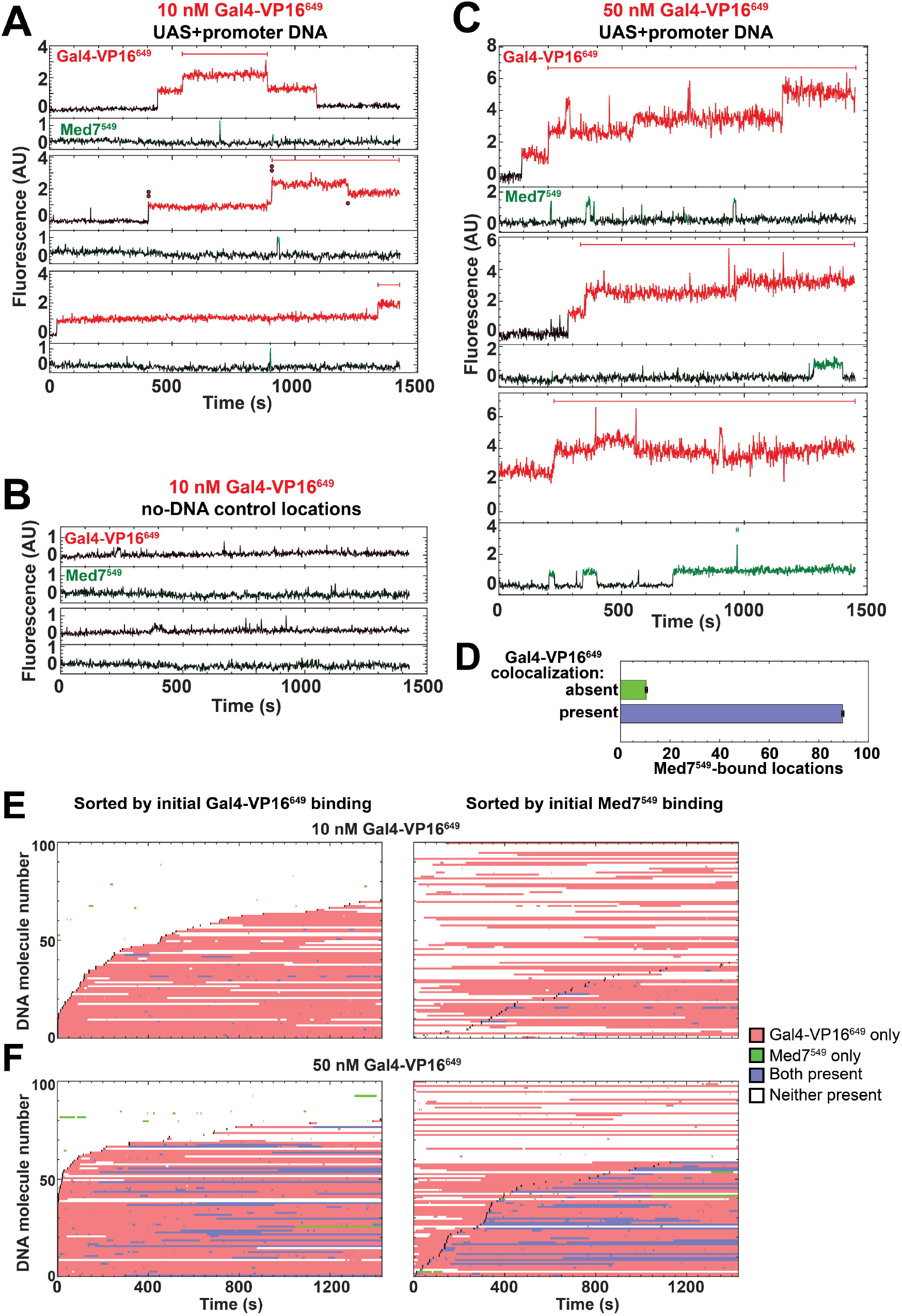
TA-dependent recruitment of Mediator to a transcription template DNA. **(A)** Example Gal4-VP16^649^ (red) and Med7^549^ (green) fluorescence intensity records from the locations of three individual DNA molecules, in YNE plus 10 nM Gal4-VP16^649^. Color: times at which a fluorescent spot was detected^**46**,**47**^. Horizontal bars: intervals during which more than one molecule was simultaneously present. Step changes in Gal4-VP16^649^ fluorescence intensity are presumed to reflect arrival or departure of dimers with either one (single dot) or two (two dots) dye molecules. **(B)** Records from two control no-DNA locations in the same experiment as (A). **(C)** Same as (A), but with 50 nM Gal4-VP16^649^. **(D)** Time-averaged fraction (± S.E.) of DNA locations with bound Mediator when Gal4-VP16^649^ either was (blue) or was not (green) simultaneously present, from the experiment in (C); *N*_DNA_ = 637 DNA molecules and *N*_t_ = 1,024 time points per molecule. **(E)** Rastergrams from 100 randomly selected DNA molecules from the experiment in (A). Each horizontal line displays the time record for a single DNA location, color-coded for Gal4-VP16^649^ and Med7^549^ presence. In the left and right plots, the same DNA molecules are ordered by the time of first Gal4-VP16^649^ binding and time of first Med7^549^ binding, respectively. The start of each first-binding event is marked (black). **(F)** Same as (E), but from experiment in (C) (50 nM Gal4-VP16^649^). See also **Figures S2** and **S3**.

Gal4 binds its DNA target site as a dimer^59^, and in some records there were two different sizes of Gal4-VP16^649^ fluorescence intensity steps (e.g., **Figure 2A**, middle trace; compare steps denoted with single and double dots), consistent with one or two active dyes. Quantitative analysis of the step increase intensity distributions (**Figure S3A**) confirmed that 97 ± 4% (S.E.) of TA dimers were labeled with at least one dye (see Methods). At a higher Gal4-VP16^649^ concentration (50 nM, **Figure 2C** and **Figure S2C**), initial TA association to a DNA molecule was commonly followed by additional stepwise increases in fluorescence intensity, indicating sequential binding of two or more TA molecules to the five Gal4 binding sites. At 50 nM activator, the progressive accumulation of TAs on individual DNAs suggests that the rate of TA binding initially outpaced the rate of departure, but eventually tended toward an equilibrium balance between arrival and departure.

While imaging TA binding, Mediator binding was simultaneously monitored. Mediator binding was DNA-specific, with little association at control no-DNA locations (**Figure 2B; Figure S2B)**. At 50nM Gal4-VP16^649^, 79% (502 of 637) of DNA locations showed Mediator binding versus only 10% (56 of 561) of control locations. At 10 nM Gal4-VP16^649^, Med7^549^ binding events were rare and short, typically lasting on the order of 1 s (e.g., **Figure 2A)**. In contrast, at 50 nM activator Mediator binding was more frequent and occupancy intervals were typically 10-to 100-fold longer (e.g., **Figure 2C**). Thus, TA concentration profoundly affected Mediator presence on the DNA.

At 50 nM activator, a single DNA usually bound only one Mediator at a time, even when multiple TA molecules were simultaneously present (e.g., **Figure 2C**, red bars). Instances of multiple Mediator molecules bound simultaneously were rare and brief (e.g., **Figure 2C**, green bar in bottom record). Multiple Mediator molecules rarely arrived simultaneously, arguing that multimolecular Mediator condensates do not form in this experimental system, as proposed for mammalian enhancers^3,41,42^.

### TA presence on DNA recruits Mediator

Our observations are consistent with a sequential recruitment model where TAs first bind DNA and then directly or indirectly bind Mediator. At 50 nM Gal4-VP16^649^, Mediator was over eight-fold more likely to occupy DNA when Gal4-VP16^649^ was present (**Figure 2D**). To rule out that Mediator arrives after TA simply because TA generally binds to DNA faster than Mediator, binding time courses for 100 randomly selected DNA molecules from the 10 nM and 50 nM Gal4-VP16^649^ experiments were plotted as two-color rastergrams (**Figure 2E, F**). Sorting by time of initial Gal4-VP16^649^ binding (left panels) shows that intervals with one or more Mediator complexes present (blue and green horizontal lines) were more frequent and longer-lived after initial TA binding than before. In contrast, when the same 100 DNA molecules were sorted by the time of initial Med7^549^ binding (**Figure 2E, F**, right, black points), TA binding intervals (e.g., **Figure 2E, F**, right, red or blue horizontal lines) were almost equally dense before and after the first Mediator binding. Furthermore, DNAs that never bound TA (**Figure 2E**, left, molecule numbers 71 – 100) almost never bound Mediator. The rare Mediator binding observed in the absence of fluorescent TA is likely explained by dark TA dimers caused by photobleaching or incomplete labeling (**Figure S3A**; see Methods) and nonspecific binding. Thus, our observations support a sequential model in which TAs bind first and remain present on the DNA to recruit Mediator.

### Mediator is recruited to UAS but stabilized by core promoter

Previous reports show Mediator interacting with TAs at the UAS^27,31,60–62^; or binding directly to the PIC at a core promoter^36,38,62,63^. To study these interactions in our system, Mediator binding to the UAS+promoter DNA was compared to “UAS-only” or “Promoter-only” constructs (**Table S4**). Two DNA constructs were sequentially attached to the same microscope slide and then simultaneously exposed to Med7^549^ YNE supplemented with unlabeled Gal4-VP16^50^ in the absence of NTPs. Since it was not imaged, we could use a higher (190 nM) concentration of unlabeled Gal4-VP16 in this experiment to increase Mediator binding. Binding was equally fast and extensive at UAS+promoter and UAS-only locations (**Figure 3A, B**). In contrast, Promoter-only DNA (identical to UAS+promoter except having the Gal4 binding sites mutated) showed much less Mediator binding (**Figure 3C, D**). This binding resembled that at control no-DNA locations (**Figure 3E**), suggesting that most binding at Promoter-only locations may reflect non-specific Mediator binding to the slide surface.

**Figure 3:**
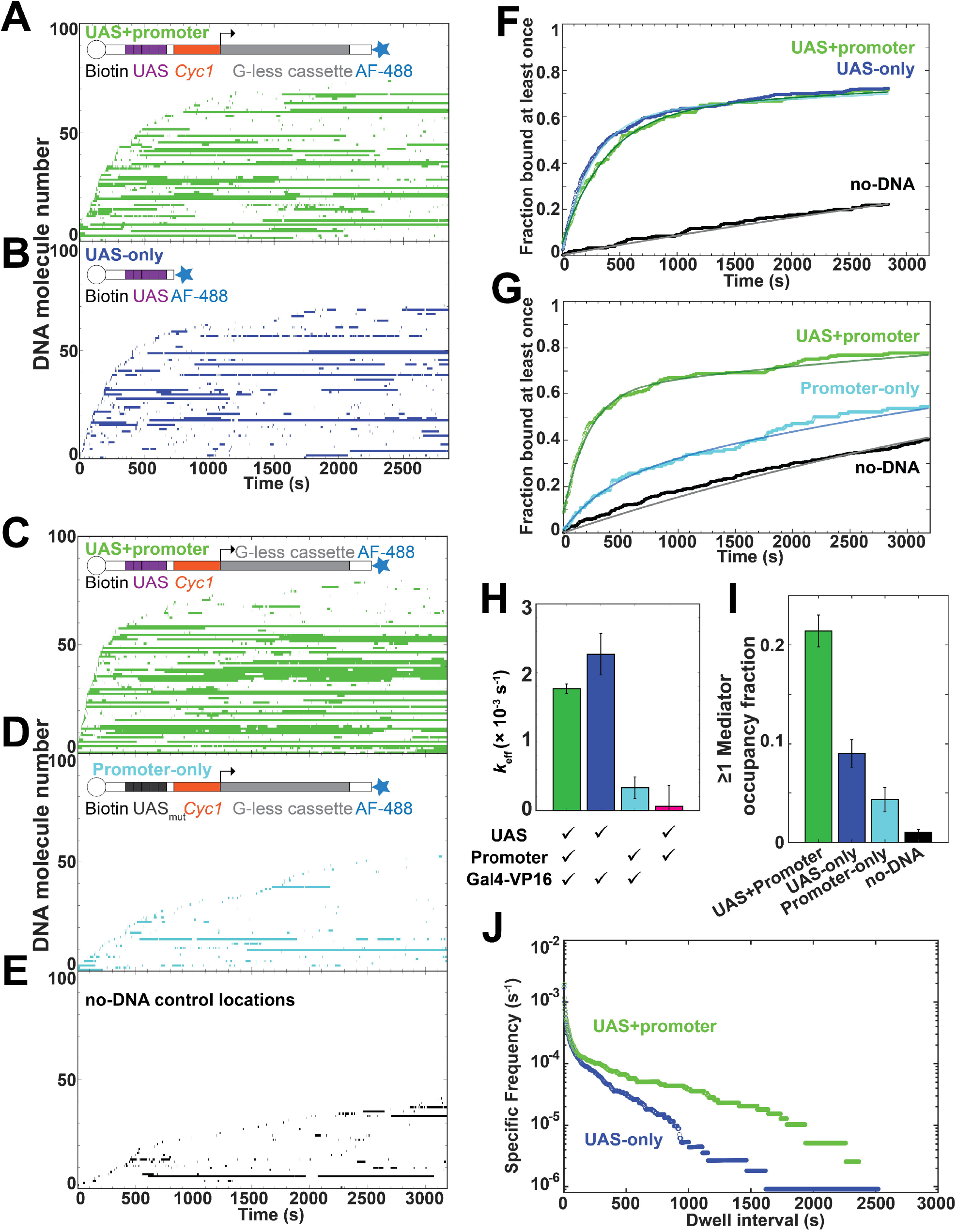
Mediator association strongly depends on UAS and activator, not core promoter. Experiments used 190 nM unlabeled Gal4-VP16. **(A, B)** Rastergrams for Med7^549^ binding at UAS+promoter DNA (A) and UAS-only DNA (B) in the same reaction (see Methods). Each plot shows data from 100 randomly selected DNA locations, sorted by the time of initial Med7^549^ arrival. Colored bars indicate times when one or more Med7^549^ molecules were bound. **(C, D)** Same as (A,B) but simultaneously monitoring UAS+promoter (C) and promoter-only (D) DNA. **(E)** Data from control no-DNA location from the same experiment as (C, D). **(F)** Cumulative fraction of UAS+promoter DNA, UAS-only DNA, and control no-DNA locations bound at least once by Med7^549^ as a function of time from addition of extract, from the experiment in (A,B). Lines are fits (see **Table S2). (G)** Same as (F), except using data from the experiment in (C-E**). (H)** Effective first-order DNA-specific association rate constants *k*_eff_ (± S.E.) for Med7^549^ measured for different DNAs with or without Gal4-VP16. Data are weighted averages of values from the replicates listed (**Table S2). (I)** Fraction of time (± S.E.) in which UAS+promoter, UAS-only, promoter-only DNAs, or no-DNA locations were bound by at least one Mediator (data pooled from Med7^549^ experiments in **Table S2**). **(J)** Cumulative distributions of Med7^549^ dwell times on UAS+promoter and UAS-only DNA in the experiment shown in (A, B). Each continuous time interval with ≥1 labeled Med7^549^ molecules present was scored as a single dwell. Specific frequency values were corrected for the contribution of non-specific surface binding at no-DNA locations (panel F, black curve). See also **Figure S4** and **Table S4**.

To quantitatively compare binding on the different DNA constructs, effective DNA-specific Mediator association rate constants (*k*_eff_, which correct for an observed slow inactivation or departure of the DNA molecules in YNE; see Methods and **Figure S4D**) were calculated using the distributions of time intervals before initial Mediator binding at each DNA or no-DNA control location^47^ (**Figure 3F, G**). The *k*_eff_ values for UAS+promoter and UAS-only DNA were identical within experimental uncertainty (**Figure 3F** and **H**, compare green and blue), suggesting that initial Mediator binding is primarily to TAs at the UAS. In agreement, DNA-specific Mediator association is essentially absent without Gal4-VP16 (**Figure 3H**, magenta; **Figure S4A-C**). Importantly, the low rate of DNA-specific Mediator binding to Promoter-only DNA suggests UAS-independent binding of Mediator directly to the core promoter makes little contribution to overall Mediator association kinetics (**Figure 3H**, compare cyan to green; **Table S2**).

Mediator association kinetics were essentially identical on UAS+promoter and UAS-only DNAs (**Figure 3F, H**). However, the fraction of time with Mediator present was substantially higher on UAS+promoter DNA (**Figure 3I**) due to longer average Mediator dwell times (**Figure 3J**). When dwell time distributions were fit to a three-component model **(Figure S4E**), the longest component was significantly lengthened by the presence of the core promoter (compare τ^L^ values in **Figure S4F**). Our previous experiments with RNApII and TFIIF (also conducted in the absence of NTPs) revealed similar recruitment by TAs to the UAS and kinetic stabilization by core promoter^49^. We propose that Mediator nucleates relatively short-lived pre-PICs containing RNApII, TFIIF, and possibly other general transcription factors^49,50,64^ while tethered to TAs at the UAS. These subsequently transfer to the core promoter sequences to form PICs, which can last for minutes in the absence of NTPs ^49,61^.

### Mediator recruits RNApII to the template

To investigate the pathway(s) by which Mediator and RNApII are assembled into the pre-PIC, we imaged fluorescently labeled Mediator (Med7^549^) and RNApII (Rpb1^Cy5^) in YNE supplemented with 190 nM unlabeled Gal4-VP16 in the absence of NTPs (**Figure 4**). Rpb1^Cy5^ bound primarily at DNA locations, with little non-specific binding at no-DNA sites (**Figure S5A-B)**. Most commonly, RNApII molecules arrived where Mediator was already present (e.g., purple-shaded intervals in **Figure 4A**). RNApII arrival without Mediator was rare (e.g., **Figure 4A**, orange; and **Figure 4B**, left), and these instances often had short dwell times resembling RNApII bound nonspecifically to the slide surface (**Figure S5B)**^50^. In contrast, Mediator generally arrived on the DNA before RNApII (**Figure 4B**), with some Mediator occupancies never recruiting RNApII (**Figure 4B**, right, DNA molecules 44-100). At this higher TA concentration, we occasionally observed two or more Med7^549^ (**Figure 4A**, green bars) or Rpb1^Cy5^ (**Figure 4A**, red bars) molecules simultaneously present on the same DNA. Simultaneously bound RNApII molecules were virtually exclusively observed at DNAs with more than one Mediator molecule.

**Figure 4:**
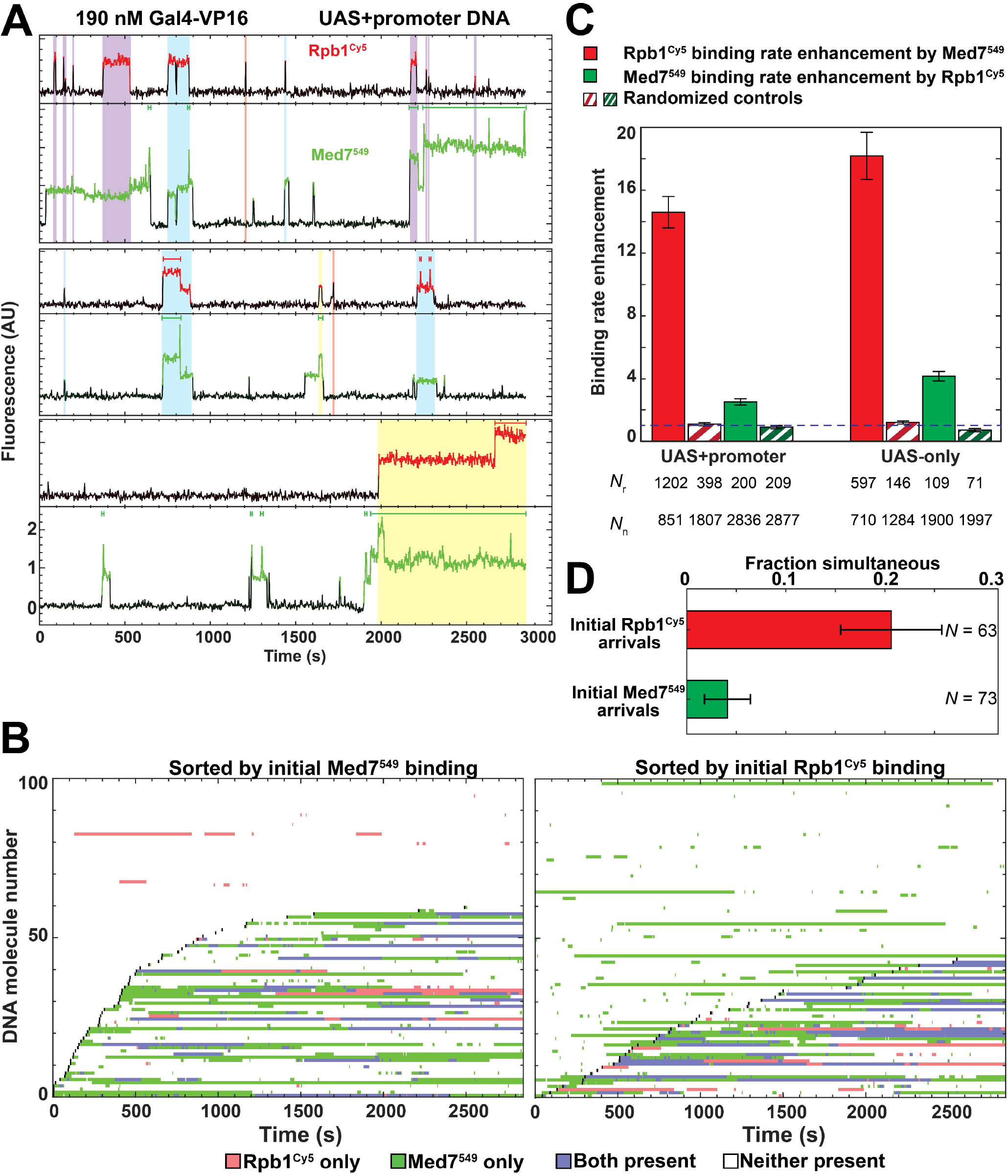
RNApII recruitment by Mediator during activator-dependent PIC assembly. **(A)** Example Rpb1^Cy5^ (red) and Med7^549^ (green) fluorescence time records recorded from the locations of three individual UAS+promoter DNA molecules (from same experiment as **Figure 3A)**. Red and green indicate times at which a fluorescent spot was detected. Red and green bars mark simultaneous presence of more than one Rpb1^Cy5^ or Med7^549^, respectively. Shading highlights RNApII-bound intervals in which RNApII arrived when Mediator was already present (purple), when both RNApII and Mediator arrived simultaneously (blue), and when RNApII arrived while no Mediator was detected (orange). Rarely observed cases in which Mediator was present before RNApII arrived simultaneously with a second Mediator molecule are shaded yellow. **(B)** Rastergrams, plotted as **Figure 2**, for 100 randomly selected DNA molecules from the experiment in (A). **(C)** Fold increase (± S.E.) in Rpb1^Cy5^ binding frequency to DNA molecules when Med7^549^ was present relative to when it was absent (red) and in Med7^549^ binding to a DNA when Rpb1^Cy5^ was present relative to when it was absent (green). Fold increase of one (dashed line) indicates no preference. *N*_r_ and *N*_n_: numbers of recruited and non-recruited binding events (see Methods). **(D)** Simultaneous arrival of the initial molecules of Mediator and RNApII to 100 randomly selected DNA locations. Red: fraction of the initial Rpb1^Cy5^ arrivals on each DNA that appeared simultaneous (within ± 2.7 s) with Med7^549^ arrival. Green: fraction of the initial Med7^549^ arrivals that appeared simultaneous with of Rpb1^Cy5^ arrival. See also **Figure S5**.

Quantitating the dependence of RNApII recruitment on Mediator, we calculated that RNApII molecules bound to Mediator-occupied UAS+promoter DNA >14-fold more frequently than without Mediator (**Figure 4C**, left, red). This enhancement was not because Mediator binds more rapidly than RNApII: a control analysis randomly pairing Mediator and RNApII records (**Figure 4C**, left, red striped bar) showed no enhancement, as expected for independent events. The enhancement of Mediator binding rate by RNApII was considerably smaller (**Figure 4C**, green bars), but greater than the randomized control value of ∼1 (**Figure 4C**, green striped bars). We attribute this small effect to occasional Med7^549^ fluorescence blinking and/or photobleaching rather than RNApII recruiting Mediator to DNA. Importantly, binding rate enhancements on UAS-only DNA (**Figure 4C**, right, red) were similar to those on UAS+promoter DNA, consistent with both Mediator and RNApII being initially recruited by UAS-bound TAs^49^.

### Mediator and RNApII can arrive at DNA in a pre-formed complex

While Mediator most often preceded RNApII on DNA, simultaneous arrivals were also common (e.g., **Figure 4A**, starts of blue and yellow intervals). Approximately 20% of initial Rpb1^Cy5^ arrivals on each DNA were simultaneous with Med7^549^ arrivals (**Figure 4D**). Simultaneous arrivals did not occur by chance alone: random time-offset control analyses exhibited zero simultaneous arrivals on 100 DNAs analyzed (see Methods). Only ∼5% of initial Mediator arrivals were accompanied by simultaneous Rpb1^Cy5^ arrivals, as expected when Mediator arrivals greatly outnumber RNApII arrivals (**Figure 4B**).

In a complementary analysis, we examined the distribution of time intervals between Mediator and RNApII arrivals on DNA (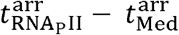 ;**Figure S5C**). The majority of ternary complexes formed with Mediator arriving prior to RNApII (**Figure S5C**, green bars), while roughly one-fifth formed by simultaneous Mediator and RNApII arrival (**Figure S5C**, blue). Kinetic analysis confirms that the apparently simultaneous arrivals cannot be accounted for by sequential arrival of the two proteins within the ± 2.7 s time resolution of the experiment (**Figure S5C**, compare blue bars to curves). Consistent with Mediator (**Figure 3**) and RNApII^49^ being initially recruited to the UAS, UAS-only results were essentially identical to UAS+promoter DNA (**Figure S5D**). Thus, some pre-formed protein complexes containing both Mediator and RNApII are present in YNE, and this fraction may be even greater in cells where protein concentrations are even higher.

### Mediator tethers RNApII to UAS-bound activator

Protein departures from DNA showed the inverse pattern of arrivals (compare **Figure S5C-D** with **E-F**). Most commonly, RNApII dissociated before Mediator. The second most common category of events was simultaneous departure. Only rarely did Mediator depart first, and this could reflect photobleaching. The arrival and departure data together are consistent with Mediator functioning as a physical tether between UAS-bound TAs and RNApII. The fact that Mediator can remain bound after RNApII dissociation raises the possibility that a single Mediator binding event at the UAS could promote multiple rounds of transcription.

### Mediator occupancy is increased synergistically by multiple activator molecules

In the simplest model for TA-Mediator interactions, one TA molecule recruits one Mediator complex. However, we noted far less Mediator occupancy at 10 nM TA than at 50 nM TA, even when a similar fraction of DNAs bound TA (compare **Figures 2E and F**). We suspected this was related to multiple TAs binding the five Gal4 sites in the UAS. Activator synergy in stimulating RNApII transcription is well-documented in vivo, but is consistent with multiple models (see Discussion)^9,11,12,14,16–19^. We therefore investigated whether cooperative recruitment of Mediator by multiple activators occurs in our system.

To quantitate the number of TA and Mediator molecules on each DNA, the Bayesian machine learning program *tapqir*^65^ was used to determine spot fluorescence intensities as a proxy for protein stoichiometry. Data from four different TA concentration experiments were analyzed (note that unlabeled TA was used for the 190 nM experiment, as high background fluorescence at that concentration would prevent single-molecule visualization). At late time points (∼1,000 s or more), individual TA and Mediator molecules were still exchanging on DNA (**Figures 2-4**), but mean fluorescence intensities of Gal4-VP16^649^ and Med7^549^ spots (**Figure S6A** and **B**, respectively) plateaued, suggesting these NTP-depleted reactions had reached equilibrium. Supporting this interpretation, the time to reach the equilibrium plateau shortened with increasing TA concentrations.

The mean number of Gal4-VP16 dimers bound at equilibrium was computed using the fractions of DNAs occupied by Gal4-VP16^649^ (**Figure S6C**, red), the fits to the Gal4-VP16^649^ intensity distribution at bound DNA molecules, (e.g., **Figure S3B**; **Supplementary data file 1**), and a correction for unlabeled Gal4-VP16 (see Methods). Gal4-VP16^649^ mean occupancy fraction (**Figure S6C**, red) increased roughly hyperbolically with concentration. Similarly, the mean number of TA molecules per DNA (**Figure 5A**, red) was well fit by a hyperbolic binding curve (Hill coefficient of 1), consistent with a simple independent binding model. Strikingly, similar analysis of Med7^549^ occupancy fraction (**Figure S6C**, green) and mean number of Mediator molecules per DNA (**Figure 5A**, green) exhibited a sigmoidal relationship with Gal4-VP16 concentration, with a marked increase from 25 to 190 nM activator. The mean number of Mediator molecules per DNA best fit the Hill equation with a coefficient of 2.9 (**Figure 5A**), consistent with switch-like or cooperative response of Mediator to TA concentration. While these results were obtained with the UAS+promoter DNA, quantitatively similar behavior was seen with the UAS-only construct (**Figure S7A**), indicating that association with the UAS and not the core promoter is the primary determinant of the occupancy curves shapes.

**Figure 5:**
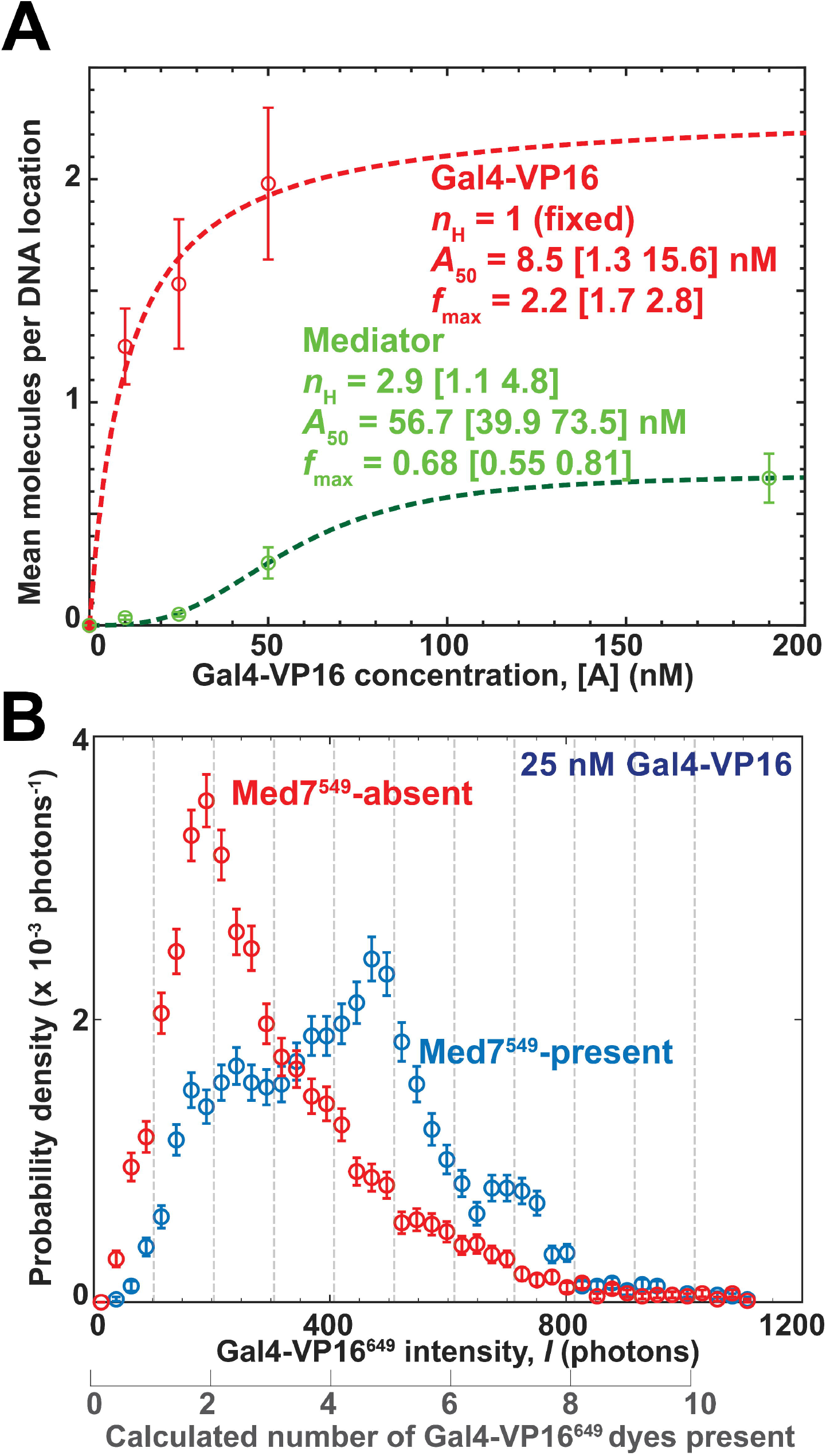
Mediator occupancy is cooperative with respect to TA concentration. (A) Mean (± S.E.) number of Gal4-VP16 and Mediator molecules per UAS+promoter DNA at equilibrium (points; see Methods and **Table S3**) at different activator concentrations [A]. Data are from the 10 and 50 nM Gal4-VP16^649^ experiments in **Figure 2**, an additional experiment at 25 nM, and the 190 nM unlabeled Gal4-VP16 experiment in **Figure 3A**. Data at zero Gal4-VP16 are zero by definition (red point) and from the experiment in **Figure S4A** (green point). Gal4-VP16 (three [A] values) and Mediator (five [A] values) were separately fit to the Hill equation 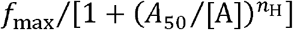 (lines) yielding the indicated parameter values [68% C.I.]. The Hill coefficient, *n*_H_ was a variable parameter for the Mediator fit but was fixed at *n*_H_ = 1 for the Gal4-VP16 fit. **(B)** Gal4-VP16^649^ equilibrium spot intensity distributions at DNA locations in the experiment at 25 nM Gal4-VP16^649^. Intensity distributions (± S.E.; *N*_DNA_ = 620) are plotted for the 3,690 data points during which Med7^549^ was present (blue points) or for a randomly-selected, time-matched set of 3,690 points during which Med7^549^ was absent (red points). Intensities are expressed both as the number of photons emitted per frame and the corresponding number of dyes present estimated from fitting intensity distributions (see Methods and **Figure S6**).

A likely explanation for this non-hyperbolic response is that DNA molecules carrying a single TA dimer interact relatively weakly with Mediator, while two or more TA present on the DNA can bind a single Mediator more strongly. To determine their relative stoichiometry, we plotted distributions of Gal4-VP16^649^ fluorescence intensities when Mediator was absent (**Figure 5B**, red points) or present (**Figure 5B**, blue points). Strikingly, Gal4-VP16^649^ fluorescence intensity on DNA without Mediator most often corresponded to two dye molecules (i.e., one activator dimer; **Figure 5B**, red points). In contrast, when Mediator was present on a DNA there was a clear shift to higher Gal4-VP16^649^ intensities corresponding to the presence of two or more TA dimers (**Figure 5B**, blue points). Very similar results were obtained using a Gal4-SNAP fusion to the yeast Gcn4 TAD (**Figure S6D**), showing that this synergy is not restricted to the VP16 TAD.

### Modeling of synergistic activator-Mediator interactions

To quantitatively understand how Mediator occupancies respond to TA concentration, we used combined TA and Mediator data (in contrast to the fits in **Figure 5A**, where TA and Mediator data were fit separately) to test model mechanisms in which up to five TA dimers bind DNA and interact with at most one Mediator molecule per DNA. For simplicity, all five possible activator-DNA contacts were assumed to have identical binding interfaces.

To start, we defined a “single-bridging” model (**Figure 6A**) allowing a single activator-Mediator interaction at a time. This model assumes TA dimers bind DNA independently and that the Mediator interacts with any single TA dimer. This model has only two variable parameters: *K*_a_, the association equilibrium constant for Gal4-VP16 binding DNA; and *m*, the strength of the Gal4-VP16−Mediator interaction (expressed as a Boltzmann factor, see Methods). Computing statistical weights associated with all molecular states produced parameter values that best fit the model to the combined equilibrium occupancy data for Gal4-VP16 and Mediator on either UAS+promoter DNA (**Figure 6B**) or UAS-only DNA (**Figure S7B**). The fits failed to reproduce the sigmoidal shape of the Mediator occupancy curves (**Figure 6B**, inset; **Figure S7B**, inset), arguing against this model.

**Figure 6:**
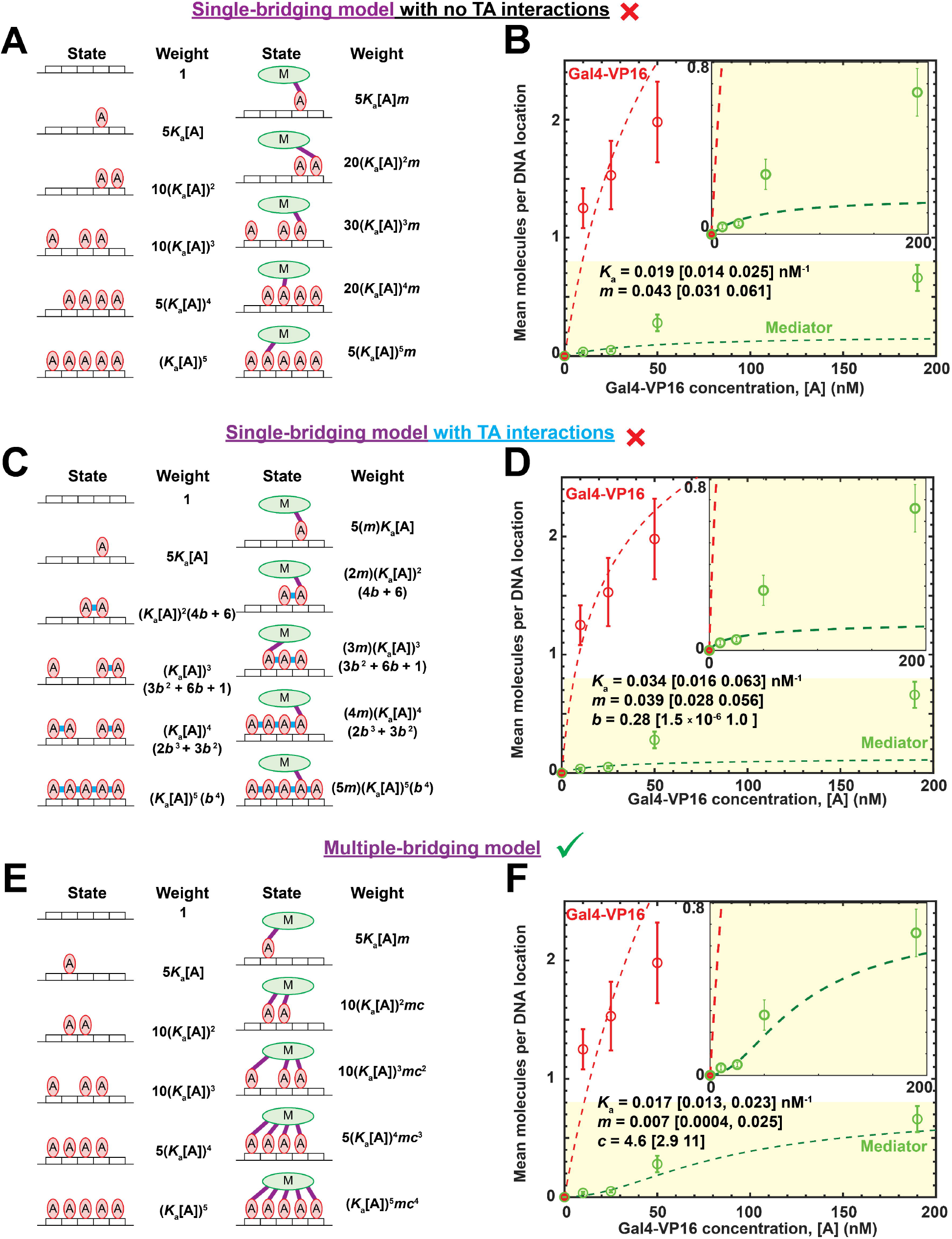
Statistical mechanical models and best fits for equilibrium TA and Mediator occupancy at the UAS. **(A)** States and statistical weights^19^ defining the single-bridging equilibrium model for binding of Gal4-VP16 dimers (red) and Mediator (green) to a DNA with five Gal4 binding sites (white rectangles). In this model, states can include a maximum of one TA-Mediator interaction (purple lines) and no TA-TA interactions. Each cartoon illustrates only one of the possible molecular arrangements included in that state; the statistical weights account for all arrangements. The weights depend on two variable parameters (*K*_a_ and *m*; see text) and the TA concentration [A]. **(B)** Mean (± S.E.) number of Gal4-VP16 and Mediator molecules per UAS+promoter DNA at equilibrium (points; same data as in **Figure 5A**). Lines are the best fit (relative likelihood *p*_6B_ = 2.06 × 10^-9^) of the **combined** Gal4-VP16 and Mediator data to the model (A), yielding the indicated parameter values [90% CI]. **(C)** Same as (A) but for the “single-bridging with TA interactions” model (*p*_6D_ = 9.71 × 10^-9^). The model is identical to that in (A) except that it adds direct interactions (with strength represented by Boltzmann factor *b*; see Methods) between adjacent TAs (blue lines). **(D)** Same data as in (B) with best fit to model in (C). **(E)** Same as (A), but for the “multiple-bridging” equilibrium model (*p*_6F_ = 2.75 × 10^-5^). The model does not include direct TA-TA interactions but assumes that every DNA-bound TA can bridge (purple) between DNA and Mediator. The weights have three variable parameters (*K*_a_, *m*, and *c*; see text). **(F)** Same data as in (B) with best fit to model in (E). Because *p*_6F_ >> *p*_6B_ or *p*_6D_, we favor the model in (E) (⍰) over those in (A) and (C) (**×**); see text and **Figure S7**.

We then considered a single-bridging model where neighboring DNA-bound TAs interact with each other (**Figure 6C**) as proposed for other activator combinations^66–69^. All the TA-TA contacts were assumed identical, with TA-TA binding energy represented by the Boltzmann factor *b*. The best fit of this model to the equilibrium occupancy data sets again failed to reproduce the shape of the Mediator occupancy curves (**Figure 6D; Figure S7C**).

We finally considered the “multiple-bridging” model shown in **Figure 6E**, which lacks direct TA-TA interactions but instead allows multiple interactions between Mediator and all TAs present on the DNA when Mediator is present. The second through fifth TA−Mediator interactions are more favorable than the first because Mediator is already in close proximity to the DNA-bound TAs, reducing the entropic cost of interaction. For simplicity, non-first TA−Mediator interactions are each assumed to have the same Boltzmann factor, *c*, so that the multiple-bridging model contains the same number of variable parameters as the single-bridging model with TA interactions. The multiple-bridging model fits the equilibrium occupancy datasets better than the single-bridging model with TA interactions (likelihood ratios *p*_6F_ / *p*_6D_ = 2.83 × 10^3^ for UAS+promoter DNA (**Figure 6D, F**) and *p*_S7D_ / *p*_S7C_ = 1.4 × 10^3^ (**Figure S7C, D**). The fits of the multiple-bridging model were in reasonable agreement with the data across all activator concentrations, reproducing the Gal4-VP16 and Mediator occupancies, the hyperbolic activator binding curve, and the sigmoidal Mediator binding curve (**Figure 6F** and **Figure S7D**). Parameter values produced by the fits roughly matched prior measurements of the interaction strengths (see Methods). Thus, a simple three-parameter model quantitatively accounted for the observed binding curves, supporting multiple TAs bridging between DNA and Mediator.

## DISCUSSION

Transcription initiation is a complex multi-step process. Here we investigate early steps in activator-dependent initiation by imaging activator, Mediator, and RNApII in the context of nuclear extract. While consistent with genomic and structural studies^39,64,70,71^, our single-molecule microscopy results provide important new insights about factor dynamics which are inaccessible using these other methods. Real-time monitoring of events on individual DNA templates reveals which PIC assembly pathways predominate (**Figure 7**, blue shading) and elucidate a mechanism by which multiple TAs could synergistically activate gene expression.

**Figure 7:**
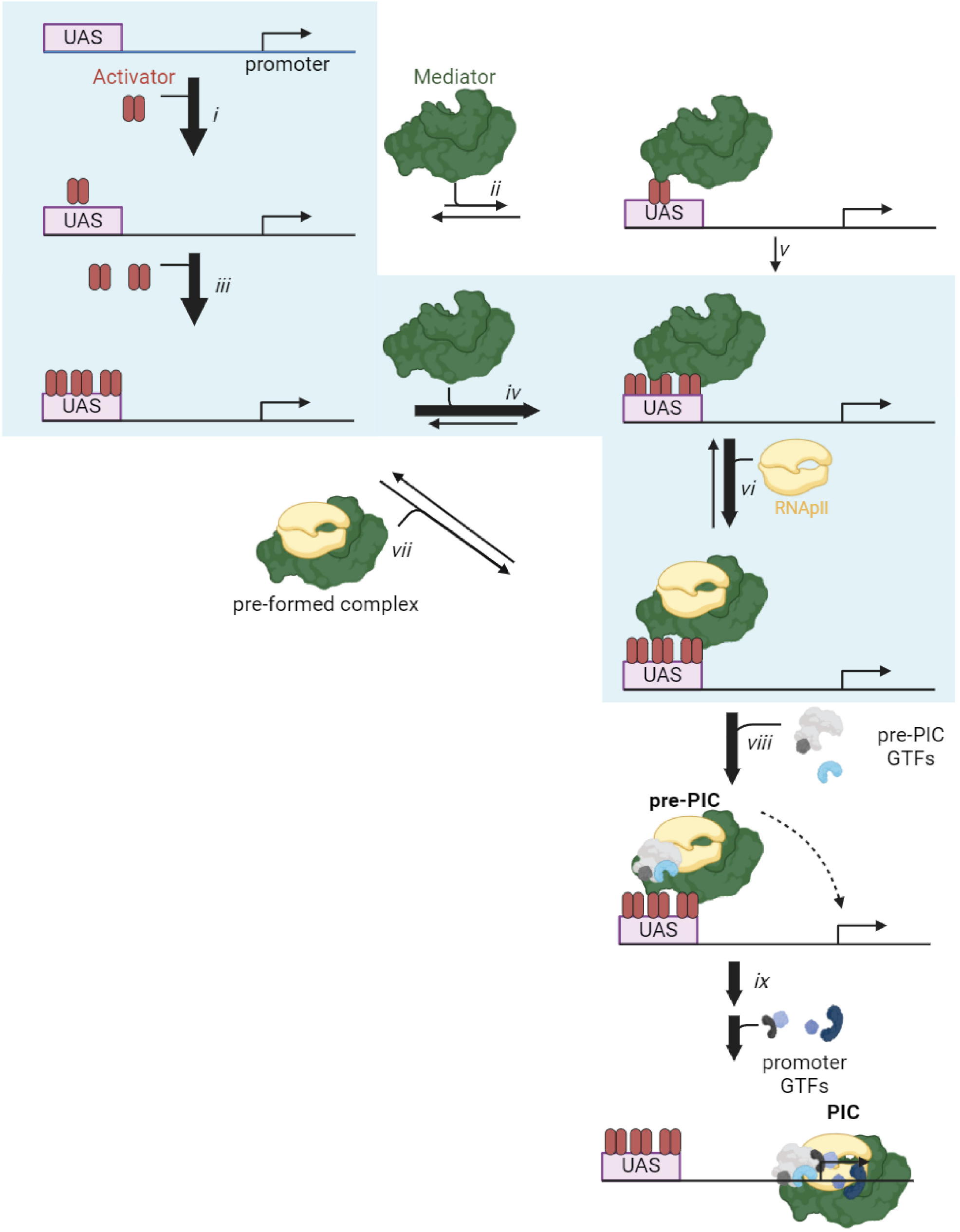
Pathways of Mediator and RNApII recruitment during activator-dependent PIC assembly. Single-molecule experiments in this study reveal hierarchical recruitment of activator, Mediator, and RNApII (depicted in red, green, and yellow, respectively) at the UAS (steps *i* through *vii*). The most commonly observed pathway for the formation of a UAS-bound pre-PIC is shaded in blue. Recruitment of Mediator to the UAS is synergistic with respect to activator concentration, likely due to multiple activator dimers tethering a single Mediator to DNA. Subsequent or simultaneous recruitment of a subset of GTFs at the UAS and transfer of the complex to the core promoter (steps *viii* and *ix*) is based on prior work^49^.

### Hierarchical interactions recruit Mediator and RNApII to the UAS

Cryo-EM structures of the PIC or initially transcribing complex (ITC) show Mediator interacting with GTFs and RNApII at the core promoter^35–37,39,72–74^. Conditional mutants show that Mediator is important for PIC assembly in vivo^63^. However, Mediator crosslinks to enhancer/UAS regions, usually more so than to core promoters, consistent with Mediator binding to UAS-associated TAs. These distinct configurations are often pictured as happening simultaneously, with Mediator creating a DNA loop between UAS-associated TAs and the promoter-associated PIC.

Our single-molecule experiments suggest a different model where TA-tethered Mediator nucleates assembly of a pre-PIC comprised of RNApII and a subset of GTFs. Initial DNA binding of Mediator requires the UAS, not the core promoter (**Figure 3**), and depends on one (**Figure 7**, steps *i, ii*) or more (**Figure 7**, steps *iii, iv*) activator molecules being on the UAS (**Figures 2, 3, S4**). This behavior mirrors RNApII and TFIIF, which we showed also initially bind at the UAS^49^. DNA molecules with multiple activator molecules showed faster Mediator association, longer Mediator dwells, and sometimes multiple Mediator molecules (**Figures 2A, C** and **4A**).

Mediator presence was required for RNApII binding to DNA (**Figure 4C**). While often modeled arriving as a pre-formed complex, we observed (**Figure 4D** and **S5C, D**) that Mediator most often arrived at the UAS before RNApII (**Figure 7**, step *vi*), and only sometimes together (**Figure 7**, step *vii*). There were many Mediator binding intervals where RNApII never appeared, but not the converse (**Figure 4B**). Therefore, RNApII-induced conformation changes in Mediator^70^ are not required for delivery to the UAS. “Empty” Mediator can bind to UAS-associated TAs and then wait at the UAS to recruit RNApII. In the absence of NTPs, RNApII left the DNA before or simultaneously with Mediator (**Figure S5E, F**), but it remains to be determined whether Mediator stays tethered to the activator after pre-PIC transfer to the core promoter.

About 20 to 35% of RNApII arrivals were simultaneous with Mediator (Figures **4D** and **S5C**,**D**), and roughly half were simultaneous with TFIIF^49^. These numbers match estimates of RNApII-Mediator and RNApII-TFIIF complexes derived from co-immunoprecipitation from extracts, although that study concluded that there was little or no trimer complex with all three factors^75^. Pre-formed Mediator-RNApII-TFIIF complexes may occur in vivo, where factor concentrations are higher. These off-DNA interactions may account for previously reported RNApII “holoenzymes”, which varied in content depending on isolation method, but characteristically contained RNApII, Mediator, and TFIIF^6,76–78^.

### Mediator pre-PICs likely transfer from UAS to core promoter

Our pre-PIC model is supported by a recent structure (designated the Med-PIC^early^ complex) showing Gal4-VP16 tethering Mediator, RNApII, TFIIF, and TFIIB at the UAS^40^. Although the intrinsically disordered VP16 TAD and Med15 Activator Binding Domains (ABDs) were not resolved, this structure is consistent with demonstrated contacts between TADs and Mediator tail subunits^30,79,80^. Tail module interactions are important for Mediator recruitment to promoters in vivo^32,81,82^, although the universality of these interactions has been disputed^28^.

Our model further proposes transfer of the pre-PIC from UAS to the core promoter (**Figure 7**, step *ix*), presumably received by TATA-binding protein. Upon PIC completion, progression to initiation is likely rapid. While transfer has not yet been directly demonstrated, several observations support this model. In the absence of NTPs to prevent transcription initiation, long Mediator dwell times increase on UAS+promoter but not UAS only DNA (**Figure 3J, Figure S4E-F**), as previously observed for RNApII and TFIIF^49^. These extended dwells likely represent PICs, as their durations are reduced by NTPs. Our results parallel in vivo ChIP experiments showing that depletion of the TFIIH kinase, which normally triggers Mediator release from the PIC, strongly increases Mediator crosslinking at the core promoter^32,61,82,83^.

### A multiple-bridging model for activator synergy in Mediator recruitment

Transcription synergy occurs when multiple activator binding sites produce a gene output greater than the sum of the individual site outputs^7,10,18,54^. Synergy is biologically significant because it can produce non-hyperbolic, switch-like responses to TA concentration and can facilitate combinatorial gene regulation by multiple TAs. A variety of mechanisms have been proposed to explain synergy^12,17^. One classic mechanism is cooperative binding of TAs at adjacent DNA sites, as experimentally observed for several prokaryotic and eukaryotic regulators^54,66–69,84^. Another model is “kinetic synergy”, where different activators accelerate distinct rate-limiting steps^9^. For example, one TA could relieve chromatin repression while a second accelerates PIC assembly.

Our results suggest a different TA synergy mechanism involving Mediator recruitment. First, the average number of bound Mediator molecules exhibited a sigmoidal dependence on TA concentration (**Figure 5A**). Second, individual DNA molecules with Mediator present were much more likely to have multiple bound TAs than DNAs without Mediator (**Figure 5B** and **Figure S6D**). The data are inconsistent with modeled single-bridging mechanisms where Mediator interacts with one TA dimer at a time, whether or not the TA dimers interact cooperatively (**Figure 6A-D**). Instead, the data are better fit by a multiple-bridging model in which two or more DNA-bound TA dimers simultaneously interact with one Mediator (**Figure 6E, F**). A generic multiple-bridging model with a single unidentified target was proposed to explain experimental observations of synergy without TA cooperativity^10,18^, and our results show that Mediator is one such target.

Activation by amphipathic TADs like VP16 and Gcn4 is strongly correlated with the strength of their binding to Med15, their principle interacting subunit on Mediator^85^. Med15 contains multiple activator-binding domains (ABDs) which can individually make dynamic, structurally pleiotropic interactions with TADs, suggesting that a single Med15 subunit may interact with multiple TADs from different activator dimers^30,86–89^, consistent with the multiple-bridging model. Our data do not exclude the possibility that some TAD-Mediator interactions might be indirect (e.g., bridging through RNApII, GTFs, or other proteins).

A single template DNA can simultaneously associate with multiple Mediators and RNApII molecules (**Figure 4A** and ref. ^49^). Thus, a cluster of TAs at the UAS may tether multiple pre-PICs, positioning them for rapid transfer to the promoter and transcript initiation. Multiple pre-PIC formation at enhancers could be a phase separation independent mechanism for the activator and Mediator clustering seen by live cell imaging. We see no evidence for formation of liquid-like condensates in our system – Gal4-VP16, Mediator, and RNApII associate with the DNA in single-molecule intensity steps^49,50^ – but when condensates are present they could provide further concentration of pre-PIC factors poised for transfer to promoters.

A multiple-bridging mechanism for Mediator recruitment by activators provides several potential advantages. First, it facilitates combinatorial regulation by requiring multiple different activators binding to the same UAS for maximal activation, creating distinct multi-input transcription responses. Even the simple compact promoters in yeast, including *HIS4*^90^, *CYC1*^91^, and *PHO5*^92^, contain multiple TA binding sites. Second, synergistic binding to multiple TAs reduces the likelihood of Mediator being competed or “squelched” by non-productive interactions with single activators molecules in the nucleoplasm. Supporting this idea, purified Mediator stably bound to DNA with multiple activators, even with a large excess of free activator molecules in solution^86^. Finally, in vivo imaging of Gal4 shows that a UAS with multiple binding sites produces transcription bursts longer than the time scale of individual Gal4 dwell times^93^.

Our studies provide new insights into the dynamics of transcription activation using a simple extract system and recombinant TAs with strong amphipathic TADs. We anticipate that such single-molecule experiments and statistical mechanical analyses will provide insight into the biochemical mechanisms of activation in systems with additional complexities such as alternative TAD types, chromatin, multiple enhancers, and long-distance enhancer-promoter communication,

### Limitations of the study

Our single-molecule studies visualize only a subset of proteins (i.e., those that are fluorescently labeled) in each experiment. Investigating the participation of other proteins or protein complexes in transcription requires additional experiments in which those proteins are labeled. Though yeast nuclear extract approximates the protein composition inside nuclei of living cells, factor concentrations are higher in vivo, leading to possible underestimation of the prevalence of protein-protein interactions. Finally, our statistical mechanical models are minimal models consistent with the data; we do not exclude the possibility that more complex models would better explain the data.

## Supporting information

Key resources table

## RESOURCE AVAILABILITY

### Lead contact

Further information and requests for resources and reagents should be directed to and will be fulfilled by the Lead Contact, Jeff Gelles.

### Materials availability

*S. cerevisiae* strains generated in this study (**Table S1**) are available from the lead contact upon request.

### Data and code availability

- Single-molecule fluorescence source data have been deposited at Zenodo and are publicly available as of the date of publication at DOI: 10.5281/zenodo.16687840.
- All original code has been deposited at Zenodo and is publicly available as of the date of publication at DOI: 10.5281/zenodo.16687840.
- Any additional information required to reanalyze the data reported in this paper is available from the lead contact upon request.

## ACKNOWLEDGEMENTS

We thank Grace Rosen and Inwha Baek for support in troubleshooting and method development, Douglas Theobald and Yerdos Ordabayev for their help with maximum likelihood modeling, Yoo Jin Joo for purified Gal4-VP16, and Johnson Chung for help with microscopy and data analysis. This work was supported by NIH grants R01CA246500 to J.G. and S.B., R01GM081648 to J.G., and R01GM046498 to S.B.

## AUTHOR CONTRIBUTIONS

J. Jeon made yeast nuclear extracts and labeled activator and performed microscopy experiments with labeled activator and Mediator on UAS+promoter DNA. N. Farheen performed microscopy experiments with labeled activator and Mediator on UAS-only DNA. D. H. Zhou performed all other experiments. Statistical mechanical modeling was performed by N. Farheen with contributions from D. H. Zhou, J. Kondev, and J. Gelles. D. H. Zhou performed microscopy image analysis with contributions from L. J. Friedman. The paper was written by D. H. Zhou, J. Jeon, S. Buratowski, and J. Gelles, with additional contributions and edits from the other authors. S. Buratowski and J. Gelles were responsible for project conception and funding acquisition.

## DECLARATION OF INTERESTS

S.B. is a member of the Molecular Cell Advisory Board. The authors declare no other competing interests.

## STAR METHODS

### EXPERIMENTAL MODEL AND STUDY PARTICIPANT DETAILS

#### *Saccharomyces cerevisiae strains* (see Table S1)

YF702 (MATa, ura3-1, leu2-3,112, trp1-1, his3-11,15, ade2-1, pep4Δ::HIS3, prbΔ::his3, prc1Δ::hisG)

YSB3499 (MATa, ura3-1, leu2-3,112, trp1-1, his3-11,15, ade2-1, pep4Δ::HIS3, prbΔ::his3, prc1Δ::hisG, RPB1-SNAPf::NATMX, MED7-HA3-DHFR::Hygromycin^R^)

YSB3613 (MATa, ura3-1, leu2-3,112, trp1-1, his3-11,15, ade2-1, pep4Δ::HIS3, prbΔ::his3, prc1Δ::hisG, RPB1-DHFR::Hygromycin^R^, MED7-HA3-SNAPf::KanMX)

YSB3687 (MATa, ura3-1, leu2-3,112, trp1-1, his3-11, 15, ade2-1, pep4Δ::HIS3, prbΔ::his3, prc1Δ::hisG, RPB1-HA3-HALO::NATMX, MED7-HA3-SNAPf::KanMX)

## METHOD DETAILS

### Purification and labeling of activator proteins

To prepare fluorescently labeled Gal4-VP16^649^, we used the Gal4-SNAP-VP16 construct as described^51^. This bacterial expression construct encodes a fusion protein consisting of the yeast Gal4 DNA binding domain (amino acids 1-95), SNAPf^94^, and the VP16 transcriptional activation domain^52^, and was subcloned in the pRJR plasmid^52^. Unlabeled Gal4-VP16 protein was prepared as in refs.^49,50^.

Gal4-SNAP-VP16 or Gal4-VP16 expression plasmids were transformed into the BL21 (codon+, DE3) *E. coli* strain. Cells were grown in LB at 37°C until the OD_600_ reached 0.4-0.5. Protein expression was then induced with 0.1 mM IPTG and 10 μM ZnCl_2_ at 30 °C for 3 hours. Cells were harvested by centrifugation at 5,000 rpm (Fiberlite F10-4x1000 LEX, Thermo Scientific) for 10 minutes at 4°C, resuspended in Lysis buffer (20 mM HEPES-KOH pH 7.6, 10 μM ZnCl_2_, 300 mM KCl, 5% (v/v) glycerol, 0.1% NP-40, and 1 mM PMSF), and lysed by sonication. The lysate was centrifuged at 15,000 rpm (Fiberlite F18-12x50, Thermo Scientific) for 15 minutes at 4°C and the supernatant was incubated with Ni-NTA agarose (Gold Biotechnology) at 4°C overnight on a rotator. The activator-bound beads were extensively washed with Wash buffer (20 mM HEPES-KOH pH 7.6, 10 μM ZnCl_2_, 30 mM KCl, 5% (v/v) glycerol, 15 mM imidazole, and 1 mM PMSF), and protein eluted with Elution buffer (20 mM HEPES-KOH pH 7.6, 10 μM ZnCl_2_, 30 mM KCl, 5% (v/v) glycerol, 600 mM imidazole, and 1 mM PMSF). Protein was further purified by salt gradient ion exchange chromatography using a Mono Q HR 5/5 column (Pharmacia) and Gradient buffers (20 mM HEPES-KOH pH 7.6, 0.1 M (low salt) or 1 M (high salt) NaCl, 10 μM ZnCl_2_, 1 mM EDTA, 20% (v/v) glycerol, 1 mM DTT, and 1 μg/ mL each of aprotinin, leupeptin, pepstatin A, and benzamidine), and dialyzed against Dialysis buffer (20 mM HEPES-KOH pH 7.6, 500 mM KOAc, 10 µM ZnCl_2_, 1 mM EDTA, and 20% (v/v) glycerol).

Purified Gal4-SNAP-VP16 was labeled with SNAP-Surface-649 (New England Biolabs) as in ref. ^51^. Approximately 0.5–5 μM protein was incubated with at least a 2-fold molar excess of dye in phosphate buffered saline (PBS) supplemented with 1 mM DTT at 4°C for 60–90 minutes. Labeled protein was incubated with Ni-NTA agarose at 4°C for 1 hour. The protein-coupled beads were extensively washed with PBS (supplemented with 10 % glycerol, 1 mM DTT, 10 μM ZnCl_2_, and 1 μg/ mL each of aprotinin, leupeptin, pepstatin A, and antipain) to remove unincorporated dye adduct. Then, labeled Gal4-VP16^649^ was eluted with PBS (supplemented with 10 % glycerol, 300 mM imidazole, 1 mM DTT, 10 μM ZnCl_2_, and 1 μg / mL each of aprotinin, leupeptin, pepstatin A, and antipain).

To construct the Gal4-SNAP-Gcn4 expression plasmid (BE573), the Gal4-SNAP-VP16 plasmid (BE559) was PCR-amplified into two fragments excluding the VP16 sequence. The Gcn4 activation domain was PCR-amplified from the BE473 plasmid using primers listed in the **Key Resources Table**. DNA fragments were ligated by isothermal assembly, and the encoded Gal4-SNAP-Gcn4 protein was purified and labeled as described for Gal4-SNAP-VP16, yielding Gal4-Gcn4^649^.

### Yeast strains and growth rate assays

*Saccharomyces cerevisiae* strains are listed in **Table S1**. Strain YSB3499 was constructed as follows. A PCR product containing the 3xHA-DHFR-Hygromycin cassette was amplified from pBS-SKII-3XHA-eDHFR-Hygromycin, YV317^49,95^ using P830 and 831 primers and transformed into the protease-deficient YF702/CB012 strain to tag the C-terminus of Med7 with HA3-DHFR. The resulting Med7-HA3-DHFR cassette, along with the hygromycin resistance marker, was then PCR-amplified from this modified strain and introduced into YSB3338 expressing Rbp1-SNAPf^49,50^ to create YSB3499. Similarly, for construction of YSB3613, a PCR product containing the 3xHA-fSNAP-KanMx cassette (from pBS-SKII-3XHA-fSNAP-Kan, YV311^95^) was amplified with the same primers and introduced into YSB3337 strain expressing Rpb1-DHFR^50,51^ to tag the C-terminus of Med7 with HA3-fSNAP, creating strain YSB3613.

To test whether the tags affect growth rates of yeast strains, yeast cultures were diluted in YPD to OD_600_ of 0.002. Multiple replicate 150 - 200 µL wells of each strain were grown with orbital agitation at 30°C for 300,000 s in a Tecan microtiter plate reader. Log phase growth rates were measured as the slope of the linear portion of the log_2_ (OD_600_) records, then normalized to the growth rate of corresponding wild-type strain (**Figure S1B**). Records without a clearly defined log phase were discarded.

### Preparation of yeast nuclear extracts

Yeast nuclear extracts were prepared as previously described^49^, with minor modifications. Yeast strains (**Table S1**) were grown in 4 L of modified YPD (1% yeast extract, 2% peptone, 3% dextrose, 0.015% tryptophan, and 0.006% adenine) at 30°C until the absorbance at 600 nm reached 3. Cells were harvested by centrifugation at 4,000 rpm (Fiberlite F10-4x1000 LEX, Thermo Scientific) for 8 minutes at 4°C and resuspended in 150 mL TD buffer (50 mM Tris-HCl pH7.5 and 30 mM DTT). After 15 minutes incubation with gentle shaking at 30°C, cells were pelleted by centrifugation at 4,000 rpm for 12 minutes at 25°C and resuspended in 20 mL YPD/S (1% yeast extract, 2% peptone, 2% dextrose, and 1 M sorbitol). We then added 15 mg of Zymolyase 100T (120493-1, Amsbio) dissolved in 30 mL of 1 M sorbitol to make spheroplasts and incubated for 30-60 min with gentle shaking at 30°C until more than 80% of cells were spheroplasts. Spheroplasts were mixed with an additional 100 mL of YPD/S and pelleted by centrifugation at 4,000 rpm for 12 minutes at 25°C. This pellet was resuspended in 250 mL YPD/S and incubated at 30°C for 30 minutes for recovery. Then spheroplasts were washed first with YPD/S and then with cold 1 M Sorbitol by sequential centrifugations at 4,000 rpm for 12 minutes at 4°C and then resuspended in 100 mL of Buffer A (18% (w/v) Ficoll 400, 10 mM Tris-HCl pH 7.5, 20 mM KOAc, 5 mM Mg(OAc)_2_, 1 mM EDTA pH 8.0, 0.5 mM spermidine, 0.17 mM spermine, 3 mM DTT, and 1 μg / mL each of aprotinin, leupeptin, pepstatin A, and antipain) and homogenized (62400-802, Wheaton). Large cell debris and improperly lysed spheroplasts were then removed as pellets by four sequential centrifugations at a low speed (twice at 5,000 rpm (Fiberlite F10-4x1000 LEX, Thermo Scientific) for 8 minutes, then twice at 5,000 rpm for 5 minutes, with new centrifuge tubes used for each spin). The final supernatant was centrifuged at 13,000 rpm (Fiberlite F10-4x1000 LEX, Thermo Scientific) for 30 minutes at 4°C to collect crude nuclei. The nuclei were resuspended in 10 mL of Buffer B (100 mM Tris-OAc pH 7.9, 50 mM KOAc, 10 mM MgSO_4_, 10% Glycerol, 2 mM EDTA pH 8.0, 3 mM DTT, and 1 μg / mL each of aprotinin, leupeptin, pepstatin A, and antipain) and lysed by addition of 3 M (NH_4_)_2_SO_4_ to a final concentration of 0.5 M. After 30 min incubation at 4°C on a rotator, the lysed nuclei were centrifuged at 37,000 rpm (Type 70 Ti, Beckman) for 90 minutes at 4°C. The nuclear proteins in the supernatant were then precipitated by a gentle addition of granular (NH_4_)_2_SO_4_ to ∼ 75 % saturation (0.35 g per 1 mL of supernatant) with 30 minutes incubation on a rotator at 4 °C. Nuclear proteins were collected by centrifugation at 13,000 rpm (type 70 Ti, Beckman) for 20 minutes, discarding the supernatant, centrifuging the pellet again for 5 minutes and again discarding supernatant. The pellet (∼0.8 g) was suspended in 2 mL of Buffer C (20 mM HEPES-KOH pH 7.6, 10 mM MgSO_4_, 1 mM EGTA pH 8.0, 10% glycerol, 3 mM DTT, and 1 μg/ mL each of aprotinin, leupeptin, pepstatin A, and antipain). SNAP-fused proteins in the extract were then labeled by incubation with 0.4 μM SNAP-Surface-549 (New England Biolabs) for 1 hour at 4°C on a rotator. The labeled nuclear extract was dialyzed against Buffer C supplemented with 75 mM (NH_4_)_2_SO_4_ or 100 mM KOAc in dialysis tubing (molecular weight cut-off, 6-8 kDa) three times (60 minutes, 90 minutes, and 120 minutes sequentially). Remaining unincorporated free dye in the nuclear extract was mostly removed by incubation with recombinant SNAPf-coupled agarose beads (Pierce 26196) at 4°C for 1 hour. The agarose beads in the nuclear extract were then removed by centrifugation at 1,000 × *g* for 2 minutes at 4 °C. The labeled nuclear extract was aliquoted, flash frozen, and stored at -80°C.

### Immunoblots

For each gel lane, 5 μL yeast nuclear extract was diluted 10-fold in 0.1% SDS, resolved by gel electrophoresis on a 10% SDS-PAGE gel, and transferred to a PVDF membrane (IPFL00010, Millipore). Anti-HA antibody (Roche, 12013819001) and anti-Rpb1 antibody (8WG16) were used for chemiluminescent detection of Med7 and Rpb1, respectively (**Figure S1C**).

### In vitro transcription activity assay

The *in vitro* bulk transcription activity assay was performed as previously described^48,49^. In brief, 250 ng SB649 plasmid^96^, 200 ng Gal4-VP16, and 10-50 µg yeast nuclear extract were incubated in ATB buffer (20 mM HEPES-KOH pH 7.6, 100 mM KOAc, 1 mM EDTA pH 8.0, and 5 mM Mg(OAc)_2_) supplemented with 10 mM phosphocreatine, 0.1-0.2 units of creatine kinase, and 0.33 units of RNAsin (Promega) at room temperature for 5 minutes. Then, 500 μM ATP, 500 μM CTP, 20 μM UTP, and 0.3 μCi of α-^32^P UTP (PerkinElmer) were added to the reaction to initiate transcription and label transcripts. After 45 minutes incubation at room temperature, transcripts were recovered by phenol/chloroform extraction, resolved by gel electrophoresis (8 M urea-6% polyacrylamide gel), and phosphoimaged on a Typhoon imager (GE Healthcare).

### DNA templates for microscope experiments

The UAS+promoter DNA template was prepared by PCR using Herculase II Fusion DNA Polymerase (Agilent #600675) from the SB649 plasmid^96^, upstream primer 5′-biotin-TTGGGTAACGCCAGGGT-3′ (IDT), and downstream primer 5′-Alexa488-AGCGGATAACAATTTCACACAG-3′. The UAS-only DNA template^49^ was prepared the same way except with downstream primer 2, 5′-Alexa488-CGAGATCCTCTAGAGTCGG-3′. The promoter-only DNA template was amplified from the SB1958 plasmid using the same primers as the UAS+promoter DNA. PCR products were purified using DNA SizeSelector-I SPRI magnetic beads (Aline Biosciences Z-6001) according to the manufacturer’s instructions. Plasmid SB1958 is identical to plasmid SB649 except that the Gal4 binding sites have been mutated. Sequences for each template are given in **Table S2**.

### Colocalization single-molecule microscopy

As previously described^49,50^, we passivated glass coverslips using mPEG-SG-2000:biotin-PEG-SVA5000 (Laysan Bio) at a 200:1 ratio, then created flow chambers by sandwiching lines of silicone vacuum grease between two coverslips. We filled the chambers with KOBS buffer (50 mM Tris-OAc (pH 8.0), 100 mM KOAc, 8 mM Mg(OAc)_2_, 27 mM NH_4_OAc, 0.1 mg/ mL bovine serum albumin (#126615 EMD Chemicals; La Jolla, CA)), and mounted them on a micro-mirror multi-wavelength TIRF microscope^46^ that used excitation wavelengths of 488 nm, 532 nm, and 633 nm. We first photobleached any fluorescent impurities on the coverslip surface by simultaneously irradiating with 2 mW at each of the three wavelengths (all laser powers were measured at the input to the micro-mirrors). We then added streptavidin-coated fluorescent beads (TransFluoSpheres, ThermoFisher Scientific T10711) to serve as fiducial markers for drift correction^46^. The slide surface was then treated with 0.013 mg/mL NeutrAvidin (ThermoFisher Scientific 31000) for one minute before being flushed with 40 µL KOBS buffer. We then introduced KOBS supplemented with ∼10 pM biotinylated and AF488-labeled DNA template, an oxygen-scavenging system (0.9 units/mL protocatechuate dioxygenase [Sigma-Aldrich P8279], 5 mM protocatechuic acid [Sigma-Aldrich 03930590]), triplet state quenchers^97^ (0.5 mM propyl gallate, 1 mM Trolox, 1 mM 4-nitrobenzyl alcohol)^98^ and 0.5% dimethylsulfoxide. Images of the DNA were acquired using the 488 nm laser at 1.2 – 1.3 mW. In experiments with multiple DNA templates (*e*.*g*., UAS+promoter and UAS-only DNA), the DNAs were sequentially added, and images were taken after only the first DNA template was attached and then after both templates were attached^99^. Finally, the reaction mix containing yeast nuclear extract, reagents, and buffers with the following final concentrations was introduced: the specified concentration of yeast nuclear extract protein, the same oxygen scavenging system and triplet state quenchers as previously detailed, NTP depletion system (20 mM glucose and 2 units hexokinase [Sigma H4502]), 20 µM Acetyl-CoA, and 20 ng / µL competitor *E. coli* genomic DNA fragments^49^, and buffer components (100.5 mM potassium acetate, 24.1 mM HEPES, pH 7.6, 0.4 mM Tris-acetate, 1 mM EDTA, 5 mM magnesium acetate, 2mM MgSO_4_, 0.2 mM EGTA, 15 mM (NH_4_)_2_SO_4_, and 4.25% glycerol). Where indicated, reactions were further supplemented with 20 nM Cy5-trimethoprim (for visualizing Rpb1-DHFR), 0 – 50 nM labeled Gal4-VP16^649^ or 190 nM unlabeled Gal4-VP16,

For experiments with Gal4-VP16^649^ (or Gal4-Gcn4^649^) and Med7^549^, separate images using 633 nm and 532 nm excitation (0.5 and 1.2 mW, respectively) were captured every 1.4 s (0.5 s for each channel, plus switching times) (**Figures 2** and **5**). For experiments with Rpb1^Cy5^ and Med7^549^, separate 633 nm and 532 nm excitation (0.4 and 0.8 mW, respectively) images were captured every 2.7 s (1 s per frame for each channel, plus switching times) (**Figures 3** and **4**; **Table S2**). Custom software Glimpse was used to operate the microscope, laser shutters, filter wheels, and camera.

### Slide surface-tethered DNA detachment/cleavage rate

To measure the rate at which surface-tethered DNA molecules detached from or were cleaved off the chamber surface (**Figure S4D**) under the conditions of the CoSMoS experiments, we first prepared chambers with surface-tethered DNA in KOBS buffer with oxygen-scavenging and triplet state quencher as described for CoSMoS experiments for DNA imaging. After imaging the DNA, we added YNE in the same reaction mix as described previously minus activator and Cy5-trimethoprim). This step was omitted for the zero-exposure control. After zero, 900, or 2,280 s exposure to yeast nuclear extract, the chamber was washed with the full CoSMoS reaction mixture (including 190 nM activator) and the DNA was immediately reimaged. The fraction of DNA molecules in the first image that had survived in the second image was normalized by the surviving fraction in the zero-exposure experiment.

### CoSMoS image analysis

Microscope images were preprocessed as previously described^47^. In brief, DNA spots were first detected in the blue-excited channel and their centers determined via two-dimensional Gaussian fitting. These locations were then mapped to the red- and green-excited “binder”^47^ protein channels and corrected for spatial drift across the time duration of the experiment using the fluorescent beads as fiducial markers. When two DNA templates were present in the same sample (e.g., both UAS+promoter and UAS-only DNAs), blue channel images pre- and post-addition of the second DNA were used to identify the two^99^. Locations lacking visible DNA molecules were also randomly selected as described^47^ to serve as controls to measure DNA-independent non-specific interactions of binder proteins with the slide surface. The fluorescence intensity values from the binder protein channels at DNA and no-DNA locations were integrated to create time records (e.g., **Figure 2A-C**; **Figure 4A**). Presence or absence of each binder protein at each DNA location in each frame was determined by applying thresholds for integrated fluorescence intensity and target-binder distance^47^. The resulting binary bound/unbound state records were used to generate rastergrams (**Figures 2E-F; 3A-E; 4B; S2; S4A-B; S5A-B**).

## QUANTIFICATION AND STATISTICAL ANALYSIS

### Mediator association rate constant

To measure DNA-specific rate constants for Mediator association (**Figure 3F-H, Table S2**), we measured the time from the beginning of the experiment until the first time a Med7^649^ molecule was detected at each DNA or no-DNA location. Restricting analysis to the first binding at each location reduces confounding effects (e.g., from photobleaching, dye blinking, or incomplete labeling) on the measurement^47^.

In the distributions of times to first binding, a minority subpopulation of DNA locations appeared to become incapable of DNA-specific Med7^649^ association, and the size of this subpopulation was lower in experiments that displayed higher DNA-specific binding rates. This observation suggests kinetic competition between DNA-specific Med7^649^ association and inhibitory processes that cause DNA locations to become refractory to Mediator association as the experiment progresses. Control experiments show that a fraction of these apparently inactive locations could be caused by DNA detachment from the surface or DNA endonucleolytic cleavage before Mediator has had a chance to bind (**Figure S4D**). However, the rate of detachment or cleavage is insufficient to account for most inactive locations. We hypothesize that most of the inactive DNAs instead result from occlusion of the DNA molecules by interactions with DNA-binding proteins (e.g., histones^48^) present in the YNE, or by non-specific interactions of the DNA molecules with the slide surface, either of which could obstruct Mediator binding. To account for these inhibitory “DNA inactivation” processes, we analyzed the time-to-first-binding data using the model

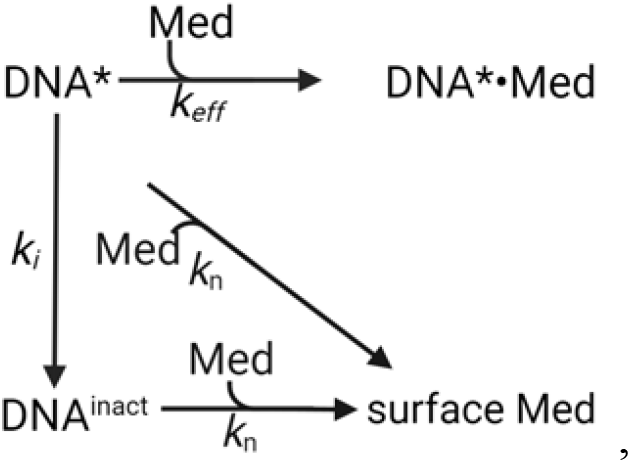

which assumes that DNA locations recorded before extract addition are initially in a state (DNA^*^) that is binding-competent, are slowly converted (at rate *k*_i_) to inactive locations DNA^inact^. Only DNA^*^ locations are capable of specific association with Mediator (Med). Specific binding (with apparent first-order rate constant *k*_eff_) forms the specific complex DNA^*^•Med. In contrast, both DNA^*^ and DNA^inact^ locations will exhibit non-specific surface binding (forming “surface Med”) with apparent first-order rate constant *k*_n_. This model predicts the conditional probability of observing a time interval *t* until first Med7^649^ binding, given specified values for the rate constants, is proportional to:

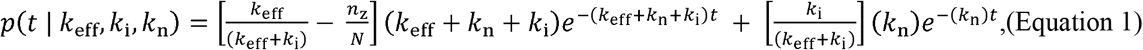

where 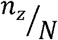 is the fraction of DNA locations at which a binder molecule was already present at the start of acquisition. To determine the rate constants *k*_eff_, *k*_i_, and *k*_n_, we adapted the model in Eqs. 4–7 of ref. ^47^, in which the three rate constants are reparametrized 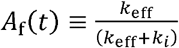 and k_on_ ≡ *k*_eff_ + *k*_i_ .We first used maximum likelihood fitting to determine *k*_n_ from data taken at control no-DNA locations. Holding *k*_n_ at this value, we then determined *A*_f_ and *k*_on_ from data at DNA locations. Finally, *k*_eff_ and *k*_i_ were calculated as *k*_eff_ *= k*_on_ × *A*_f_ and *k*_i_ = *k*_on_ (1 - *A*_f_). Parameter values with S.E. calculated by bootstrapping (1,000 samples) are reported in **Table S2**. Values of *k*_i_ could only be accurately determined in experimental conditions (UAS-containing DNA and Gal4-VP16 present) that showed specific Med7^649^ binding to DNA. In those experiments the value of *k*_i_ was consistently around 1–2 × 10^-3^ s^-1^ (**Table S2**), consistent with the model assumption of a constant rate of time-dependent DNA inactivation.

### Mediator dwell times

Frequency distributions of Med7^649^ specific-binding dwell times on DNA (**Figure 3J**) were determined by subtracting the distribution measured at randomly selected no-DNA locations from the distribution at DNA locations measured from the same recording^47^. Only dwells for which the end of the dwell was observed during the experiment were used in this analysis. Distributions were plotted as cumulative dwell time frequency distributions or as binned dwell time probability density distributions.

Measured dwell times were fit to a multi-exponential model, using an maximum likelihood approach analogous to that described by Equations 9 – 12 of ref. ^47^ (**Figure S4E-F**). First, dwells from non-specific Mediator binding to the slide surface at no-DNA locations were fit using the empirical biexponential probability density model

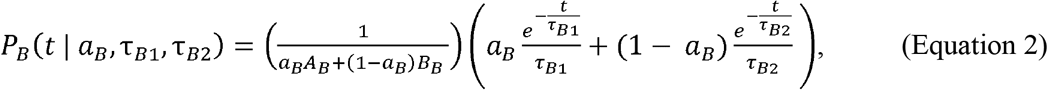

with 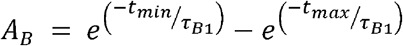 and 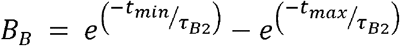. The variables *t*_min_ and *t*_max_ were set to the minimum and maximum interval lengths that can be resolved in the experiment. The two exponential components have characteristic lifetimes τ_B1_ and τ_B2_ and amplitudes *a*_B_ and (1 − *a*_B_). The corresponding likelihood function was maximized to obtain values for *a*_B,_ τ_B1_, and τ_B2_.

Next, dwells from DNA locations were fit to triexponential empirical probability density model for DNA-specific Mediator binding that also includes the biexponential non-specific surface binding determined from no-DNA locations:

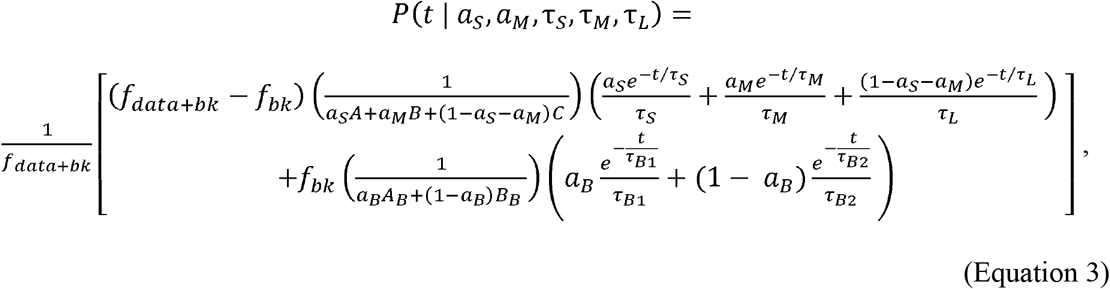

with 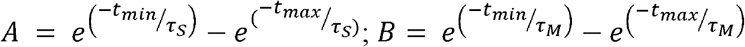 and 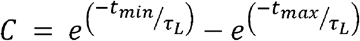 .The variables *f*_*data+bk*_ and *f*_*bk*_ were set to the frequencies of Mediator binding events measured at DNA and no-DNA locations, respectively. The three exponential components have characteristic lifetimes τ_S_, τ_M_, and τ_L_ and amplitudes *a*_S_, *a*_M_, and *a*_L_ = (1 −*a*_*S*_ − *a*_*M*_). This corresponding likelihood equation maximized to determine values for the parameters τ_S_, τ_M_, and τ_L_, *a*_S_, and *a*_M_.

### Binding rate enhancement

To measure the acceleration by the presence of Mediator on the DNA binding of RNApII, we counted the number of times in a single-molecule experiment that RNApII arrived at a DNA location at which Mediator was already bound, *N*_Med->Med-RNApII_ ≡ *N*_r_, and the analogous number of RNApII arrivals when Mediator was not already bound, *N*_ø->RNApII_ ≡ *N*_n_. We then calculated the recruited and non-recruited RNApII binding frequencies as *f*_r_ = *N*_r_ / *t*_r_ and *f*_n_ = *N*_n_ / *t*_n_, respectively, where *t*_r_ is the number of time points at which Mediator, but not RNApII, was present and *t*_n_ is the number of time points at which neither Mediator nor RNApII was present. The subset of binding events in which Med7^549^ and Rpb1^Cy5^ arrived within 2.7 s of each other were excluded from this analysis. We also calculated an analogous quantity *f*_b_ = *N*_b_ / *t*_b_ for control no-DNA locations to account for the non-specific background binding of Mediator. The DNA-specific binding rate enhancement (**Figure 4C**, solid red bars) was taken to be

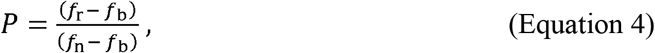

where *P* = 1 indicates equal preference for the two binding modes. To check that P > 1 values were not coincidental, we also performed randomized control analyses in which Mediator and RNApII records from different DNA locations were randomly selected without replacement and paired for the *P* calculation. These controls yielded *P* ≈ 1 (**Figure 4C**, red striped bars) confirming the detected preferences were not biased by kinetic differences between the proteins studied. These steps were repeated to analyze binding rate enhancement of Mediator binding by RNApII (**Figure 4C**, solid green and green striped bars). In **Figure 4C** left, the binding rate enhancement values are calculated from DNA-specific RNApII binding frequencies of Med7^549^ and Rpb1^Cy5^ to UAS+promoter DNA in the experiment in **Figure 4A** plus two additional replicates. In **Figure 4C** right they are calculated from a single replicate experiment with UAS-only DNA.

The standard error in P was calculated as

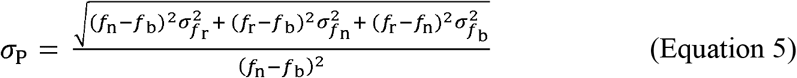

where the standard errors in respective binding frequencies are 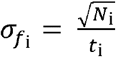.

### Simultaneous arrival of Mediator and RNApII

Mediator and RNApII were scored as arriving simultaneously (**Figure 4D**; **Figure S5C-D**) if appearance of Med7^549^ and Rpb1^Cy5^ fluorescence at a DNA location occurred within the experimental time resolution (± 2.7 s). Control analyses of the same 100 randomly selected first-binding datasets in **Figure 4D** in which a random time offset (uniformly distributed between −133 and +133 seconds) was introduced between the Rpb1^Cy5^ and Med7^549^ record from each DNA location exhibited zero simultaneous arrivals, indicating that the observed simultaneous arrivals of Med7^549^ and Rpb1^Cy5^ were not coincidental.

To test the idea that apparently simultaneous arrivals of Med7^549^ and Rpb1^Cy5^ reflected recruitment of a pre-formed complex containing both proteins, rather than independent sequential binding of the two proteins within the time resolution of the experiment, we quantitatively examined the formation of Mediator and RNApII ternary complexes. Ternary complexes were defined using time intervals in which both Med7^549^ and Rpb1^Cy5^ were simultaneously present for at least one shared frame. To minimize the effect of Cy5-trimethoprim blinking, only a single Rpb1^Cy5^ event was considered per Med7^549^ dwell (for example, see **Figure 4A**, top trace between 0 – 600 seconds). Ternary complex formation events were then categorized as in **Figure S5C-D** (bars). The kinetically modeled sequential events (red and green fit curves) were substantially lower than the blue bars when extrapolated to zero time-difference, indicating that the vast majority of these apparently simultaneous arrivals were indeed simultaneous. The analogous analysis was performed for simultaneous departures (**Figure S5E-F**).

### Spot presence and spot fluorescence intensity

CoSMoS experiments with Med7^549^ and either Gal4-VP16^649^, Gal4-VP16, or Gal4-Gcn4^649^ were processed by the Bayesian machine learning method implemented in the software *tapqir* using the described time-independent *cosmos* model^65^. To measure the fraction of DNA molecules colocalized with a Gal4-VP16^649^ or Med7^549^ fluorescence spot in these experiments (**Figure S6C**), we scored time points at each DNA location with a DNA-specific Gal4-VP16^649^ and/or Med7^549^ spot as those with specific spot probability *p*_specific_ ≥ 0.5. In the experiment with no Gal4-VP16^649^, the fraction of DNAs with a colocalized Gal4-VP16^649^ spot was taken to be zero (leading to the solid red points in **Figure 5A; Figure 6B, D, F;** and **Figure S6C**).

The *cosmos* model fits two candidate spots per frame per DNA location. For both candidates it calculates the probabilities *p*(θ = 1) and *p*(θ = 2) (denoted *theta_probs* in ref. ^100^) that candidate spots 1 and 2 are authentic, target-specific spots^65^. To accommodate cases in which a single, DNA-colocalized spot was fit as consisting of two overlapping candidate spots, we take the intensity (in photons) above background of all target-specific (i.e., *p*_specific_ ≥ 0.5) spots to be

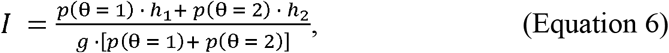

where the *cosmos* model parameters *h*_1_ and *h*_2_ are the integrated Gaussian-modeled spot intensities for spots 1 and 2; and *g* is camera gain (ADU/photon). This theta probability weighting means the intensity value will reflect the “true” spot when the choice is clear or combine the signals when a single intense spot is modeled as two overlapping spots. To calculate the time-dependence of the average Gal4-VP16^649^ and Med7^549^ fluorescence per DNA molecule (**Figure S6A, B**, respectively), we binned the experimental timepoints into quintiles (10, 25, 50 nM Gal4-VP16) or deciles (190 nM Gal4-VP16; see **Table S3**). For all DNA locations at all timepoints in each bin, we averaged the spot intensity *I* (Equation 6) if a target-specific spot was detected (i.e., *p*_specific_ ≥ 0.5) or a value of zero if no spot was detected (*p*_specific_ < 0.5). Standard errors of this mean were conservatively estimated based on the number of DNA molecules, *N*_DNA_ (i.e., they did not account for the number of time points), since successive measurements on the same DNA location are not fully statistically independent.

### Fitting spot intensity distributions

The molecular complexes studied may in general contain more than one Gal4-VP16 or Mediator and therefore may contain more than one dye-labeled Gal4-VP16^649^ or Med7^549^ protein. To measure the fluorescence intensities, distributions of target-specific fluorescence spot intensity *I* across multiple time points and DNA locations (e.g., **Figure S3)** were maximum likelihood fit to the gamma distribution mixture model probability density function

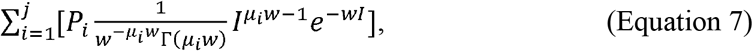

where *j* was set at a fixed number of components corresponding to the maximum number of dyes per spot; *µ*_*i*_ was taken to be *i* ·*µ*_1_ for *i* = 2,…,*j*; and the fit parameters were *µ*_1_, the mean intensity above background of a single dye moiety; *w*, the gamma distribution inverse scale parameter; and *P*_*i*_ for *i* = 1,…,*j*, the fractional amplitude of the *i*^th^ component, where 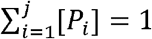. All fits, fit parameter values, and parameter standard errors are reported in **Supplementary Data File 1**.

### Gal4-VP16^649^ and Med7^549^ fraction labeled

The fraction of Gal4-VP16 dimers in the Gal4-VP16^649^ preparation that contained one or more fluorescent dye moieties was measured from the distribution of fluorescence intensity changes accompanying the first Gal4-VP16^649^ binding seen at each DNA location in the 10 nM Gal4-VP16^649^ experiment (**Figure 2A**). In the first instance of three consecutive Gal4-VP16^649^-present (*p*_specific_ ≥ 0.5) frames immediately following a Gal4-VP16^649^-absent (*p*_specific_ < 0.5) frame, we used the intensity above background *I* (Equation 6) values for the second *p*_specific_ ≥ 0.5 frame. The distribution of these intensities was fit to Equation 7 as described above, using *j* = 4 (**Figure S3A)**. Gal4-VP16 binds tightly to its cognate DNA sequence as a dimer, so we assume that the first and second gamma components correspond to dimers with one or two fluorescent dyes, respectively, while the much smaller amplitude third and fourth gamma components are due to outlier events, e.g., instances in which two dimers bound in rapid succession. The subunit labeling fraction was calculated as

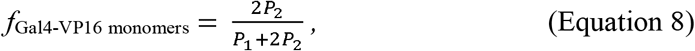

and the fraction of dimers with at least one dye was then calculated as

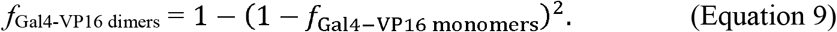

The reported experiments used a single Gal4-VP16^649^ preparation with *f*_Gal4-VP16 monomers_ = 82 ± 11% and *f*_Gal4-VP16 dimers_=97±4%.

We estimated a lower limit on the fraction of dye-labeled Mediator molecules in the Med7^549^/Rpb1^Cy5^ extract from the ratio of background-corrected frequencies for Rpb1^Cy5^ arrival when Med7^549^ was present to when it was absent intervals, which is the 14.6 ± 1.0 recruitment preference (**Figure 4C**, left, red bar). This corresponds to a lower limit Med^549^ labeling fraction of 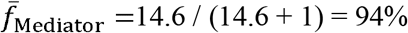.

### Number of Gal4-VP16 or Med7 molecules per DNA

To calculate the average number of Gal4-VP16 or Med7 on each DNA at equilibrium (**Figure 5A**, points; **Figure 6**, points), *I* (Equation 6) distributions were fit to Equation 7, using values of *j* chosen to account for the observed distributions while using the smallest number of components to prevent overparameterization: Gal4-VP16^649^ intensities were fit using *j* = 8 or 10; and Med7^549^ intensities were fit with *j* = 4 (**Supplementary Data File 1**). For Gal4-VP16^649^, equilibrium *µ*_1_ values were similar at all concentrations and were similar to *µ*_1_ measured earlier from the first binding event data (compare **Figure S3A** with **Figure S3B** and **Supplementary Data File 1)**, consistent with the fitting procedure accurately measuring the single-dye intensity values in the equilibrium intensity distributions. The *µ*_1_ values for the equilibrium Med7^549^ intensity distributions from the experiments with 10, 25, and 50 nM Gal4-VP16^649^ were also consistent with each other, but not with data at 190 nM activator, which was collected under different acquisition conditions as described earlier. At zero Gal4-VP16^649^, there was not enough Med7^549^ binding to fit an intensity distribution to the Gamma mixture model, so all instances of binding were assumed to have one Med7^549^ molecule present. We calculated the average number of dye molecules per fluorescent spot as

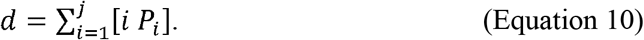

The fraction of uncleaved UAS+promoter DNA molecules remaining on the slide surface during the equilibrium time intervals, each of which was centered at *t*_c_ = ∼1,250 s from the start of the experiment, was estimated as 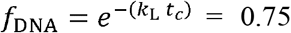, where *k*_L_ = 2.3 × 10^-4^ s^-1^ is the independently determined rate of DNA loss from the surface (**Figure S4D**). Analogous values for UAS-only DNA experiments (**Figure S7**) are reported in **Supplementary Data File 1**. The mean numbers of Gal4-VP16 dimers and Mediator complexes per surviving DNA location were then calculated as

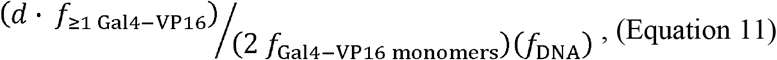

and

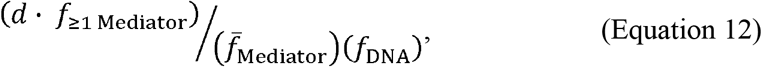

respectively, where *f* _≥ 1Gal − VP16_ *f* _≥ 1 Mediator_ are the fractions of original DNA locations with one or more dye-labeled Gal4-VP16^649^ and Mediator^549^ (**Figure 5A**).

### Quantitative models for Mediator recruitment

To test quantitative explanations for the observed activator binding and Mediator recruitment by activator to a DNA containing five identical Gal4 binding sites, we formulated three contrasting equilibrium statistical mechanics models: two “single-bridging” models with either noninteracting or interacting TA molecules (**Figures 6A** and **C**, respectively) and a “multiple-bridging” model (**Figure 6E)**. For each model, we enumerated all possible protein-bound states of a DNA molecule and calculated the associated statistical weights from the free energy contributions of the protein-DNA and protein-protein interactions present and the state multiplicity^19^. The probability of a DNA being in a given state at equilibrium is the state statistical weight divided by the sum of all weights. In each model, the state probabilities depend on the free concentration of Gal4-VP16, [A] (which is assumed to be equal to the total added Gal4-VP16 or Gal4-VP16^649^), and either two or three variable parameters.

In the independent single-bridging model (**Figure 6A**), in which only one TA at a time can interact with Mediator, only two parameters were needed: *K*_a_, the equilibrium constant for association of a TA dimer with a single Gal4 binding site on the DNA; and *m* = [M]*K*_m_, where [M] is the concentration of Mediator and *K*_m_ is the equilibrium constant for association of a single Mediator to a single TA dimer on DNA.

In the single-bridging model with TA interactions (**Figure 6C**), an additional parameter was introduced to reflect the interactions between TAs: 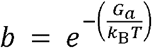, the Boltzmann factor representing the strength of each TA interaction, where *G*_a_ is the free energy associated with interactions between a pair of neighboring TAs bound to a DNA.

For the multiple-bridging model (**Figure 6E**), state probabilities depend on the model parameters *K*_*a*_, *m*, and additional parameter *c*. The parameter 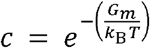 is the Boltzmann factor corresponding to the free energy *G*_m_ associated with the interaction between each of the second through fifth TAs and Mediator.

To determine the set of parameter values, 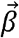, that give the best agreement between a model and the occupancy data, we used the MATLAB *fminsearch* function to minimize:

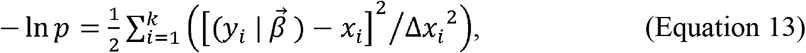

where *p* is the relative likelihood, *x*_i_ are the experimental measurements of mean numbers of activator dimers or Mediator molecules per DNA (the points shown for non-zero activator concentrations in **Figure 6** and **S7**), Δ*x*_*i*_ is the standard error associated with each *x*_i_, and 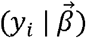 are the corresponding model predictions given the set of model parameters. *k* = 7 and *k* = 14 are the number of measurements in the data sets for the UAS+promoter (**Figure 6**) and UAS-only (**Figure S7**) DNAs, respectively. Parameter confidence intervals were determined by bootstrapping (1,000 samples).

For the multiple-bridging model the value of *K*_a_ determined from the fit (0.017 nM^-1^) is of similar magnitude to a published measurement (0.067 nM^-1^; ref. ^89^) for Gal4-DBD binding to its consensus DNA site at a somewhat different ionic strength. The fitting procedure estimates *m*, the product of Mediator concentration and its association constant with a single DNA-bound activator. Since single molecules cannot be detected by TIRF above solution concentrations of roughly 50 nM, the concentration of Mediator in our experiments is at or below that value. Thus, the fit value of *m*, 0.007, yields a predicted single activator-Mediator association equilibrium constant of *K*_m_ ≥ 0.14 µM^-1^, in rough agreement with equilibrium constants reported for binding of Mediator to Gal4 and Gcn4 TADs^29,30,88^.

## Supplemental information

Document S1. Figures S1–S7, Tables S1–S4, and supplemental references

Supplementary Data File 1: Fits of spot intensity distributions to 10- and 4-component Gamma mixture models, related to Figure 6, Figure S3, and Figure S7.

## Notes

### Competing Interest Statement

The authors have declared no competing interest.

### Summary of Updates

Reorganized and updated including new Figures S6D and S7.

## References

1. Reményi, A., Schöler, H.R., and Wilmanns, M. (2004). Combinatorial control of gene expression. Nat. Struct. Mol. Biol. 11, 812–815. 10.1038/nsmb820.

2. Weake, V.M., and Workman, J.L. (2010). Inducible gene expression: diverse regulatory mechanisms. Nat. Rev. Genet. 11, 426–437. 10.1038/nrg2781.

3. Cramer, P. (2019). Organization and regulation of gene transcription. Nature 573, 45–54. 10.1038/s41586-019-1517-4.

4. Struhl, K. (1991). Mechanisms for diversity in gene expression patterns. Neuron 7, 177–181. 10.1016/0896-6273(91)90256-Y.

5. Sainsbury, S., Bernecky, C., and Cramer, P. (2015). Structural basis of transcription initiation by RNA polymerase II. Nat. Rev. Mol. Cell Biol. 16, 129–143. 10.1038/nrm3952.

6. Lee, T.I., and Young, R.A. (2000). Transcription of Eukaryotic Protein-Coding Genes. Annu. Rev. Genet. 34, 77–137. 10.1146/annurev.genet.34.1.77.

7. Ptashne, M., and Gann, A. (1997). Transcriptional activation by recruitment. Nature 386, 569–577. 10.1038/386569a0.

8. Ptashne, M. (2005). Regulation of transcription: from lambda to eukaryotes. Trends Biochem. Sci. 30, 275–279. 10.1016/j.tibs.2005.04.003.

9. Herschlag, D., and Johnson, F.B. (1993). Synergism in transcriptional activation: a kinetic view. Genes Dev. 7, 173–179. 10.1101/gad.7.2.173.

10. Lin, Y.-S., Carey, M., Ptashne, M., and Green, M.R. (1990). How different eukaryotic transcriptional activators can cooperate promiscuously. Nature 345, 359–361. 10.1038/345359a0.

11. Carey, M. (1998). The Enhanceosome and Transcriptional Synergy. Cell 92, 5–8. 10.1016/S0092-8674(00)80893-4.

12. Kim, S., and Wysocka, J. (2023). Deciphering the multi-scale, quantitative cis-regulatory code. Mol. Cell 83, 373–392. 10.1016/j.molcel.2022.12.032.

13. Sharon, E., Kalma, Y., Sharp, A., Raveh-Sadka, T., Levo, M., Zeevi, D., Keren, L., Yakhini, Z., Weinberger, A., and Segal, E. (2012). Inferring gene regulatory logic from high-throughput measurements of thousands of systematically designed promoters. Nat. Biotechnol. 30, 521–530. 10.1038/nbt.2205.

14. Spitz, F., and Furlong, E.E.M. (2012). Transcription factors: from enhancer binding to developmental control. Nat. Rev. Genet. 13, 613–626. 10.1038/nrg3207.

15. Giorgetti, L., Siggers, T., Tiana, G., Caprara, G., Notarbartolo, S., Corona, T., Pasparakis, M., Milani, P., Bulyk, M.L., and Natoli, G. (2010). Noncooperative Interactions between Transcription Factors and Clustered DNA Binding Sites Enable Graded Transcriptional Responses to Environmental Inputs. Mol. Cell 37, 418–428. 10.1016/j.molcel.2010.01.016.

16. Scholes, C., DePace, A.H., and Sánchez, Á. (2017). Combinatorial Gene Regulation through Kinetic Control of the Transcription Cycle. Cell Syst. 4, 97-108.e9. 10.1016/j.cels.2016.11.012.

17. Reiter, F., Wienerroither, S., and Stark, A. (2017). Combinatorial function of transcription factors and cofactors. Curr. Opin. Genet. Dev. 43, 73–81. 10.1016/j.gde.2016.12.007.

18. Carey, M., Lin, Y.S., Green, M.R., and Ptashne, M. (1990). A mechanism for synergistic activation of a mammalian gene by GAL4 derivatives. Nature 345, 361–364. 10.1038/345361a0.

19. Bintu, L., Buchler, N.E., Garcia, H.G., Gerland, U., Hwa, T., Kondev, J., and Phillips, R. (2005). Transcriptional regulation by the numbers: models. Curr. Opin. Genet. Dev. 15, 116–124. https://doi.org/16/j.gde.2005.02.007.

20. Bhoite, L.T., Yu, Y., and Stillman, D.J. (2001). The Swi5 activator recruits the Mediator complex to the HO promoter without RNA polymerase II. Genes Dev. 15, 2457–2469. 10.1101/gad.921601.

21. Bryant, G.O., and Ptashne, M. (2003). Independent Recruitment In Vivo by Gal4 of Two Complexes Required for Transcription. Mol. Cell 11, 1301–1309. 10.1016/S1097-2765(03)00144-8.

22. Holstege, F.C.P., Jennings, E.G., Wyrick, J.J., Lee, T.I., Hengartner, C.J., Green, M.R., Golub, T.R., Lander, E.S., and Young, R.A. (1998). Dissecting the Regulatory Circuitry of a Eukaryotic Genome. Cell 95, 717–728. 10.1016/S0092-8674(00)81641-4.

23. Thompson, C.M., and Young, R.A. (1995). General requirement for RNA polymerase II holoenzymes in vivo. Proc. Natl. Acad. Sci. U. S. A. 92, 4587–4590.

24. Richter, W.F., Nayak, S., Iwasa, J., and Taatjes, D.J. (2022). The Mediator complex as a master regulator of transcription by RNA polymerase II. Nat. Rev. Mol. Cell Biol. 23, 732–749. 10.1038/s41580-022-00498-3.

25. Chen, H., and Pugh, B.F. (2021). What do Transcription Factors Interact With? J. Mol. Biol. 433, 166883. 10.1016/j.jmb.2021.166883.

26. Poss, Z.C., Ebmeier, C.C., and Taatjes, D.J. (2013). The Mediator complex and transcription regulation. Crit. Rev. Biochem. Mol. Biol. 48, 575–608. 10.3109/10409238.2013.840259.

27. Allen, B.L., and Taatjes, D.J. (2015). The Mediator complex: a central integrator of transcription. Nat. Rev. Mol. Cell Biol. 16, 155–166. 10.1038/nrm3951.

28. Warfield, L., Donczew, R., Mahendrawada, L., and Hahn, S. (2022). Yeast Mediator facilitates transcription initiation at most promoters via a Tail-independent mechanism. Mol. Cell 82, 4033-4048.e7. 10.1016/j.molcel.2022.09.016.

29. Herbig, E., Warfield, L., Fish, L., Fishburn, J., Knutson, B.A., Moorefield, B., Pacheco, D., and Hahn, S. (2010). Mechanism of Mediator Recruitment by Tandem Gcn4 Activation Domains and Three Gal11 Activator-Binding Domains. Mol. Cell. Biol. 30, 2376–2390. 10.1128/MCB.01046-09.

30. Tuttle, L.M., Pacheco, D., Warfield, L., Wilburn, D.B., Hahn, S., and Klevit, R.E. (2021). Mediator subunit Med15 dictates the conserved “fuzzy” binding mechanism of yeast transcription activators Gal4 and Gcn4. Nat. Commun. 12, 2220. 10.1038/s41467-021-22441-4.

31. Sun, F., Sun, T., Kronenberg, M., Tan, X., Huang, C., and Carey, M.F. (2021). The Pol II preinitiation complex (PIC) influences Mediator binding but not promoter–enhancer looping. Genes Dev. 35, 1175–1189. 10.1101/gad.348471.121.

32. Knoll, E.R., Zhu, Z.I., Sarkar, D., Landsman, D., and Morse, R.H. (2018). Role of the pre-initiation complex in Mediator recruitment and dynamics. eLife 7, e39633. 10.7554/eLife.39633.

33. Petrenko, N., Jin, Y., Wong, K.H., and Struhl, K. (2016). Mediator Undergoes a Compositional Change during Transcriptional Activation. Mol. Cell 64, 443–454. 10.1016/j.molcel.2016.09.015.

34. Zhao, H., Li, J., Xiang, Y., Malik, S., Vartak, S.V., Veronezi, G.M.B., Young, N., Riney, M., Kalchschmidt, J., Conte, A., et al. (2024). An IDR-dependent mechanism for nuclear receptor control of Mediator interaction with RNA polymerase II. Mol. Cell 84, 2648-2664.e10. 10.1016/j.molcel.2024.06.006.

35. Robinson, P.J., Trnka, M.J., Bushnell, D.A., Davis, R.E., Mattei, P.-J., Burlingame, A.L., and Kornberg, R.D. (2016). Structure of a Complete Mediator-RNA Polymerase II Pre-Initiation Complex. Cell 166, 1411-1422.e16. 10.1016/j.cell.2016.08.050.

36. Plaschka, C., Larivière, L., Wenzeck, L., Seizl, M., Hemann, M., Tegunov, D., Petrotchenko, E.V., Borchers, C.H., Baumeister, W., Herzog, F., et al. (2015). Architecture of the RNA polymerase II–Mediator core initiation complex. Nature 518, 376–380. 10.1038/nature14229.

37. Schilbach, S., Hantsche, M., Tegunov, D., Dienemann, C., Wigge, C., Urlaub, H., and Cramer, P. (2017). Structures of transcription pre-initiation complex with TFIIH and Mediator. Nature 551, 204–209. 10.1038/nature24282.

38. Plaschka, C., Nozawa, K., and Cramer, P. (2016). Mediator Architecture and RNA Polymerase II Interaction. J. Mol. Biol. 428, 2569–2574. 10.1016/j.jmb.2016.01.028.

39. Rengachari, S., Schilbach, S., Aibara, S., Dienemann, C., and Cramer, P. (2021). Structure of the human Mediator–RNA polymerase II pre-initiation complex. Nature 594, 129–133. 10.1038/s41586-021-03555-7.

40. Gorbea Colón, J.J., Palao, L., III, Chen, S.-F., Kim, H.J., Snyder, L., Chang, Y.-W., Tsai, K.-L., and Murakami, K. (2023). Structural basis of a transcription pre-initiation complex on a divergent promoter. Mol. Cell 83, 574-588.e11. 10.1016/j.molcel.2023.01.011.

41. Sabari, B.R., Dall’Agnese, A., Boija, A., Klein, I.A., Coffey, E.L., Shrinivas, K., Abraham, B.J., Hannett, N.M., Zamudio, A.V., Manteiga, J.C., et al. (2018). Coactivator condensation at super-enhancers links phase separation and gene control. Science 361, eaar3958. 10.1126/science.aar3958.

42. Cho, W.-K., Spille, J.-H., Hecht, M., Lee, C., Li, C., Grube, V., and Cisse, I.I. (2018). Mediator and RNA polymerase II clusters associate in transcription-dependent condensates. Science 361, 412. 10.1126/science.aar4199.

43. Roeder, R.G. (2019). 50+ years of eukaryotic transcription: an expanding universe of factors and mechanisms. Nat. Struct. Mol. Biol. 26, 783–791. 10.1038/s41594-019-0287-x.

44. Kornberg, R.D. (2005). Mediator and the mechanism of transcriptional activation. Trends Biochem. Sci. 30, 235–239. 10.1016/j.tibs.2005.03.011.

45. Harper, T.M., and Taatjes, D.J. (2018). The complex structure and function of Mediator. J. Biol. Chem. 293, 13778–13785. 10.1074/jbc.R117.794438.

46. Friedman, L.J., Chung, J., and Gelles, J. (2006). Viewing Dynamic Assembly of Molecular Complexes by Multi-Wavelength Single-Molecule Fluorescence. Biophys. J. 91, 1023–1031. 10.1529/biophysj.106.084004.

47. Friedman, L.J., and Gelles, J. (2015). Multi-wavelength single-molecule fluorescence analysis of transcription mechanisms. Methods 86, 27–36. 10.1016/j.ymeth.2015.05.026.

48. Sikorski, T.W., Joo, Y.J., Ficarro, S.B., Askenazi, M., Buratowski, S., and Marto, J.A. (2012). Proteomic Analysis Demonstrates Activator- and Chromatin-specific Recruitment to Promoters. J. Biol. Chem. 287, 35397–35408. 10.1074/jbc.M112.391581.

49. Baek, I., Friedman, L.J., Gelles, J., and Buratowski, S. (2021). Single-molecule studies reveal branched pathways for activator-dependent assembly of RNA polymerase II pre-initiation complexes. Mol. Cell 81, 3576-3588.e6. 10.1016/j.molcel.2021.07.025.

50. Rosen, G.A., Baek, I., Friedman, L.J., Joo, Y.J., Buratowski, S., and Gelles, J. (2020). Dynamics of RNA polymerase II and elongation factor Spt4/5 recruitment during activator-dependent transcription. Proc. Natl. Acad. Sci. 117, 32348–32357. 10.1073/pnas.2011224117.

51. Jeon, J., Friedman, L.J., Zhou, D.H., Seo, H.D., Adeleke, O.A., Graham, B., Patteson, E.F., Gelles, J., and Buratowski, S. (2025). Single-molecule analysis of transcription activation: dynamics of SAGA coactivator recruitment. Nat. Struct. Mol. Biol. 32, 675–686. 10.1038/s41594-024-01451-y.

52. Sadowski, I., Ma, J., Triezenberg, S., and Ptashne, M. (1988). GAL4-VP16 is an unusually potent transcriptional activator. Nature 335, 563.

53. Croston, G.E., Laybourn, P.J., Paranjape, S.M., and Kadonaga, J.T. (1992). Mechanism of transcriptional antirepression by GAL4-VP16. Genes Dev. 6, 2270–2281. 10.1101/gad.6.12a.2270.

54. Giniger, E., and Ptashne, M. (1988). Cooperative DNA binding of the yeast transcriptional activator GAL4. Proc. Natl. Acad. Sci. U. S. A. 85, 382–386. 10.1073/pnas.85.2.382.

55. Joo, Y.J., Ficarro, S.B., Chun, Y., Marto, J.A., and Buratowski, S. (2019). In vitro analysis of RNA polymerase II elongation complex dynamics. Genes Dev. 33, 578–589. 10.1101/gad.324202.119.

56. Joo, Y.J., Ficarro, S.B., Soares, L.M., Chun, Y., Marto, J.A., and Buratowski, S. (2017). Downstream promoter interactions of TFIID TAFs facilitate transcription reinitiation. Genes Dev. 31, 2162–2174. 10.1101/gad.306324.117.

57. Griggs, D.W., and Johnston, M. (1991). Regulated expression of the GAL4 activator gene in yeast provides a sensitive genetic switch for glucose repression. Proc. Natl. Acad. Sci. U. S. A. 88, 8597–8601. 10.1073/pnas.88.19.8597.

58. Jiang, F., Frey, B.R., Evans, M.L., Friel, J.C., and Hopper, J.E. (2009). Gene Activation by Dissociation of an Inhibitor from a Transcriptional Activation Domain. Mol. Cell. Biol. 29, 5604–5610. 10.1128/MCB.00632-09.

59. Hong, M., Fitzgerald, M.X., Harper, S., Luo, C., Speicher, D.W., and Marmorstein, R. (2008). Structural Basis for Dimerization in DNA Recognition by Gal4. Structure 16, 1019–1026. 10.1016/j.str.2008.03.015.

60. Ebmeier, C.C., and Taatjes, D.J. (2010). Activator-Mediator binding regulates Mediator-cofactor interactions. Proc. Natl. Acad. Sci. 107, 11283–11288. 10.1073/pnas.0914215107.

61. Jeronimo, C., and Robert, F. (2014). Kin28 regulates the transient association of Mediator with core promoters. Nat. Struct. Mol. Biol. 21, 449–455. 10.1038/nsmb.2810.

62. Sikorski, T.W., and Buratowski, S. (2009). The basal initiation machinery: beyond the general transcription factors. Curr. Opin. Cell Biol. 21, 344–351. 10.1016/j.ceb.2009.03.006.

63. Jean-Jacques, H., Poh, S.L., and Kuras, L. (2018). Mediator, known as a coactivator, can act in transcription initiation in an activator-independent manner in vivo. Biochim. Biophys. Acta BBA - Gene Regul. Mech. 1861, 687–696. 10.1016/j.bbagrm.2018.07.001.

64. Bernecky, C., Grob, P., Ebmeier, C.C., Nogales, E., and Taatjes, D.J. (2011). Molecular Architecture of the Human Mediator–RNA Polymerase II–TFIIF Assembly. PLOS Biol. 9, e1000603. 10.1371/journal.pbio.1000603.

65. Ordabayev, Y.A., Friedman, L.J., Gelles, J., and Theobald, D.L. (2022). Bayesian machine learning analysis of single-molecule fluorescence colocalization images. eLife 11, e73860. 10.7554/eLife.73860.

66. Adams, C.C., and Workman, J.L. (1995). Binding of disparate transcriptional activators to nucleosomal DNA is inherently cooperative. Mol. Cell. Biol. 15, 1405–1421. 10.1128/MCB.15.3.1405.

67. Kang, T., Martins, T., and Sadowski, I. (1993). Wild type GAL4 binds cooperatively to the GAL1-10 UASG in vitro. J. Biol. Chem. 268, 9629–9635.

68. Weinberg, R.L., Veprintsev, D.B., Bycroft, M., and Fersht, A.R. (2005). Comparative binding of p53 to its promoter and DNA recognition elements. J. Mol. Biol. 348, 589–596. 10.1016/j.jmb.2005.03.014.

69. Malecka, K.A., Ho, W.C., and Marmorstein, R. (2009). Crystal Structure of a p53 Core Tetramer Bound to DNA. Oncogene 28, 325–333. 10.1038/onc.2008.400.

70. Soutourina, J. (2018). Transcription regulation by the Mediator complex. Nat. Rev. Mol. Cell Biol. 19, 262–274. 10.1038/nrm.2017.115.

71. Tsai, K.-L., Yu, X., Gopalan, S., Chao, T.-C., Zhang, Y., Florens, L., Washburn, M.P., Murakami, K., Conaway, R.C., Conaway, J.W., et al. (2017). Mediator structure and rearrangements required for holoenzyme formation. Nature 544, 196–201. 10.1038/nature21393.

72. Plaschka, C., Hantsche, M., Dienemann, C., Burzinski, C., Plitzko, J., and Cramer, P. (2016). Transcription initiation complex structures elucidate DNA opening. Nature 533, 353–358. 10.1038/nature17990.

73. Hantsche, M., and Cramer, P. (2017). Conserved RNA polymerase II initiation complex structure. Curr. Opin. Struct. Biol. 47, 17–22. 10.1016/j.sbi.2017.03.013.

74. Nozawa, K., Schneider, T.R., and Cramer, P. (2017). Core Mediator structure at 3.4 Å extends model of transcription initiation complex. Nature 545, 248–251. 10.1038/nature22328.

75. Rani, P.G., Ranish, J.A., and Hahn, S. (2004). RNA polymerase II (Pol II)-TFIIF and Pol II-mediator complexes: the major stable Pol II complexes and their activity in transcription initiation and reinitiation. Mol. Cell. Biol. 24, 1709–1720. 10.1128/MCB.24.4.1709-1720.2004.

76. Kim, Y.-J., Björklund, S., Li, Y., Sayre, M.H., and Kornberg, R.D. (1994). A multiprotein mediator of transcriptional activation and its interaction with the C-terminal repeat domain of RNA polymerase II. Cell 77, 599–608. 10.1016/0092-8674(94)90221-6.

77. Koleske, A.J., and Young, R.A. (1994). An RNA polymerase II holoenzyme responsive to activators. Nature 368, 466–469. 10.1038/368466a0.

78. Koleske, A.J., Chao, D.M., and Young, R.A. (1996). Purification of yeast RNA polymerase II holoenzymes. Methods Enzymol. 273, 176–184. 10.1016/S0076-6879(96)73018-5.

79. Tuttle, L.M., Pacheco, D., Warfield, L., Luo, J., Ranish, J., Hahn, S., and Klevit, R.E. (2018). Gcn4-Mediator Specificity Is Mediated by a Large and Dynamic Fuzzy Protein-Protein Complex. Cell Rep. 22, 3251–3264. 10.1016/j.celrep.2018.02.097.

80. Zhang, F., Sumibcay, L., Hinnebusch, A.G., and Swanson, M.J. (2004). A triad of subunits from the Gal11/tail domain of Srb mediator is an in vivo target of transcriptional activator Gcn4p. Mol. Cell. Biol. 24, 6871–6886. 10.1128/MCB.24.15.6871-6886.2004.

81. Ansari, S.A., and Morse, R.H. (2012). Selective role of Mediator tail module in the transcription of highly regulated genes in yeast. Transcription 3, 110–114. 10.4161/trns.19840.

82. Jeronimo, C., Langelier, M.-F., Bataille, A.R., Pascal, J.M., Pugh, B.F., and Robert, F. (2016). Tail and Kinase Modules Differently Regulate Core Mediator Recruitment and Function In Vivo. Mol. Cell 64, 455–466. 10.1016/j.molcel.2016.09.002.

83. Wong, K.H., Jin, Y., and Struhl, K. (2014). TFIIH Phosphorylation of the Pol II CTD Stimulates Mediator Dissociation from the Preinitiation Complex and Promoter Escape. Mol. Cell 54, 601–612. 10.1016/j.molcel.2014.03.024.

84. Ptashne, M. (2014). The Chemistry of Regulation of Genes and Other Things. J. Biol. Chem. 289, 5417–5435. 10.1074/jbc.X114.547323.

85. Sanborn, A.L., Yeh, B.T., Feigerle, J.T., Hao, C.V., Townshend, R.J.L., Lieberman-Aiden, E., Dror, R.O., and Kornberg, R.D. (2021). Simple biochemical features underlie transcriptional activation domain diversity and dynamic, fuzzy binding to Mediator. eLife 10, e68068. 10.7554/eLife.68068.

86. Sanborn, A.L., Yeh, B.T., Feigerle, J.T., Hao, C.V., Townshend, R.J., Lieberman Aiden, E., Dror, R.O., and Kornberg, R.D. (2021). Simple biochemical features underlie transcriptional activation domain diversity and dynamic, fuzzy binding to Mediator. eLife 10, e68068. 10.7554/eLife.68068.

87. Hahn, S., and Young, E.T. (2011). Transcriptional regulation in Saccharomyces cerevisiae: transcription factor regulation and function, mechanisms of initiation, and roles of activators and coactivators. Genetics 189, 705–736. 10.1534/genetics.111.127019.

88. Brzovic, P.S., Heikaus, C.C., Kisselev, L., Vernon, R., Herbig, E., Pacheco, D., Warfield, L., Littlefield, P., Baker, D., Klevit, R.E., et al. (2011). The acidic transcription activator Gcn4 binds the mediator subunit Gal11/Med15 using a simple protein interface forming a fuzzy complex. Mol. Cell 44, 942–953. 10.1016/j.molcel.2011.11.008.

89. Wands, A.M., Wang, N., Lum, J.K., Hsieh, J., Fierke, C.A., and Mapp, A.K. (2011). Transient-state kinetic analysis of transcriptional activator·DNA complexes interacting with a key coactivator. J. Biol. Chem. 286, 16238–16245. 10.1074/jbc.M110.207589.

90. Tice-Baldwin, K., Fink, G.R., and Arndt, K.T. (1989). BAS1 Has a Myb Motif and Activates HIS4 Transcription Only in Combination with BAS2. Science 246, 931–935. 10.1126/science.2683089.

91. Guarente, L., Lalonde, B., Gifford, P., and Alani, E. (1984). Distinctly regulated tandem upstream activation sites mediate catabolite repression of the CYC1 gene of S. cerevisiae. Cell 36, 503–511. 10.1016/0092-8674(84)90243-5.

92. Vogel, K., Hörz, W., and Hinnen, A. (1989). The two positively acting regulatory proteins PHO2 and PHO4 physically interact with PHO5 upstream activation regions. Mol. Cell. Biol. 9, 2050–2057. 10.1128/mcb.9.5.2050-2057.1989.

93. Pomp, W., Meeussen, J.V.W., and Lenstra, T.L. (2024). Transcription factor exchange enables prolonged transcriptional bursts. Mol. Cell 84, 1036-1048.e9. 10.1016/j.molcel.2024.01.020.

94. Keppler, A., Gendreizig, S., Gronemeyer, T., Pick, H., Vogel, H., and Johnsson, K. (2003). A general method for the covalent labeling of fusion proteins with small molecules in vivo. Nat. Biotechnol. 21, 86–89. 10.1038/nbt765.

95. Baek, I., Le, S.N., Jeon, J., Chun, Y., Reed, C., and Buratowski, S. (2022). A set of Saccharomyces cerevisiae integration vectors for fluorescent dye labeling of proteins. G3 Bethesda 12, jkac201. 10.1093/g3journal/jkac201.

96. Johnson, A., Li, G., Sikorski, T.W., Buratowski, S., Woodcock, C.L., and Moazed, D. (2009). Reconstitution of Heterochromatin-Dependent Transcriptional Gene Silencing. Mol. Cell 35, 769–781. 10.1016/j.molcel.2009.07.030.

97. Dave, R., Terry, D.S., Munro, J.B., and Blanchard, S.C. (2009). Mitigating Unwanted Photophysical Processes for Improved Single-Molecule Fluorescence Imaging. Biophys. J. 96, 2371–2381. 10.1016/j.bpj.2008.11.061.

98. Crawford, D.J., Hoskins, A.A., Friedman, L.J., Gelles, J., and Moore, M.J. (2007). Visualizing the splicing of single pre-mRNA molecules in whole cell extract. RNA 14, 170–179. 10.1261/rna.794808.

99. Friedman, L.J., Mumm, J.P., and Gelles, J. (2013). RNA polymerase approaches its promoter without long-range sliding along DNA. Proc. Natl. Acad. Sci. 110, 9740–9745. 10.1073/pnas.1300221110.

100. Tapqir Documentation (2022). https://tapqir.readthedocs.io/en/stable/. https://tapqir.readthedocs.io/en/stable/.

101. Thompson, N.E., Aronson, D.B., and Burgess, R.R. (1990). Purification of eukaryotic RNA polymerase II by immunoaffinity chromatography. Elution of active enzyme with protein stabilizing agents from a polyol-responsive monoclonal antibody. J. Biol. Chem. 265, 7069–7077.

102. Hoskins, A.A., Friedman, L.J., Gallagher, S.S., Crawford, D.J., Anderson, E.G., Wombacher, R., Ramirez, N., Cornish, V.W., Gelles, J., and Moore, M.J. (2011). Ordered and Dynamic Assembly of Single Spliceosomes. Science 331, 1289–1295. 10.1126/science.1198830.

103. Inada, T., Winstall, E., Tarun, S.Z., Yates, J.R., Schieltz, D., and Sachs, A.B. (2002). One-step affinity purification of the yeast ribosome and its associated proteins and mRNAs. RNA 8, 948–958.

104. Reeves, W.M., and Hahn, S. (2003). Activator-Independent Functions of the Yeast Mediator Sin4 Complex in Preinitiation Complex Formation and Transcription Reinitiation. Mol. Cell. Biol. 23, 349–358. 10.1128/MCB.23.1.349-358.2003.

105. Jeon, J., Friedman, L.J., Zhou, D.H., Seo, H.D., Adeleke, O.A., Graham, B., Patteson, E.F., Gelles, J., and Buratowski, S. (2025). Single-molecule analysis of transcription activation: dynamics of SAGA coactivator recruitment. Nat. Struct. Mol. Biol. 32, 675–686. 10.1038/s41594-024-01451-y.

106. Cho, E.-J., Takagi, T., Moore, C.R., and Buratowski, S. (1997). mRNA capping enzyme is recruited to the transcription complex by phosphorylation of the RNA polymerase II carboxy-terminal domain. Genes Dev. 11, 3319–3326. 10.1101/gad.11.24.3319.

107. Johnson, A., Li, G., Sikorski, T.W., Buratowski, S., Woodcock, C.L., and Moazed, D. (2009). Reconstitution of Heterochromatin-Dependent Transcriptional Gene Silencing. Mol. Cell 35, 769–781. 10.1016/j.molcel.2009.07.030.

108. Baek, I., Friedman, L.J., Gelles, J., and Buratowski, S. (2021). Single-molecule studies reveal branched pathways for activator-dependent assembly of RNA polymerase II pre-initiation complexes. Mol. Cell 81, 3576–3588. 10.1016/j.molcel.2021.07.025.

109. Friedman, L.J. (2015). CoSMoS_Analysis - Tools for analyzing CoSMoS image data. https://github.com/gelles-brandeis/CoSMoS_Analysis.

110. Ordabayev, Y.A., Friedman, L.J., Gelles, J., and Theobald, D.L. (2022). Bayesian machine learning analysis of single-molecule fluorescence colocalization images. eLife 11, e73860. 10.7554/eLife.73860.

