## Supplementary material for "A mechanism of synergistic Mediator recruitment in RNA polymerase II transcription activation revealed by single-molecule fluorescence": Key resources table

| REAGENT or RESOURCE | SOURCE | IDENTIFIER |
| --- | --- | --- |
| Antibodies | | |
| Monoclonal anti-HA, peroxidase conjugated (3F10) | Roche | Cat#12013819001; RRID: AB_390917 |
| Monoclonal anti-Rpb1 CTD (8WG16) | Thompson et al.^101^ | N/A |
| Bacterial and virus strains | | |
| BL21(codon+, DE3) | Stratagene | Cat#230245 |
| Chemicals, peptides, and recombinant proteins | | |
| IPTG | Gold Biotechnology | Cat#I2481C25 |
| Ni-NTA agarose | Gold Biotechnology | Cat#PI88222 |
| Mono Q HR 5/5 column | Pharmacia | Cat#17-0546-01 |
| PMSF | Gold Biotechnology | Cat#P-470-25 |
| Aprotinin | Gold Biotechnology | Cat#A-655-100 |
| Leupeptin | Biosynth International | Cat#ILP-4041 |
| Pepstatin A | Biosynth International | Cat#IPA-4397 |
| Benzamidine | Sigma-Aldrich | Cat#B6506 |
| Antipain | Biosynth International | Cat#IAP-4062 |
| SNAP-Surface 649 | New England Biolabs | Cat#S9159S |
| SNAP-Surface 549 | New England Biolabs | Cat#S9112S |
| Cy5-TMP | Hoskins et al.^102^ | N/A |
| Yeast extract | VWR | Cat#76348-890 |
| Peptone | VWR | Cat#76347-602 |
| Dextrose | VWR | Cat#BT213090 |
| Tryptophan | Sigma-Aldrich | Cat#T0254 |
| Adenine | Sigma-Aldrich | Cat#A9126 |
| Sorbitol | VWR | Cat#97062-206  Cat#76348-156 |
| Zymolyase 100T | Amsbio | Cat#120493-1 |
| Ficoll 400 | Accurate Chemical and Scientific Corp. | Cat#AN-228520 |
| Spermidine trihydrochloride | Sigma-Aldrich | Cat#S2501 |
| Spermine tetrahydrochloride | VWR | Cat#IC10047401 |
| DTT | Oakwood Products, Inc. | Cat#M02712 |
| Ammonium sulfate | VWR | Cat#EM-AX1385-3 |
| Phosphocreatine | Sigma-Aldrich | Cat#P7936 |
| Creatine kinase | Sigma-Aldrich | Cat#C3755 |
| RNasin | Promega | Cat#N2511 |
| α-32P UTP | PerkinElmer | Cat#BLU-507H |
| mPEG-SG-2000 | Laysan Bio | Cat#MPEG-SG-2000-1GR |
| Biotin-PEG-SVA5000 | Laysan Bio | Cat#BIO-PEG-SVA-5K/mPEG-SVA-5K |
| Bovine serum albumin | EMD Chemicals | Cat#126615 |
| TransFluoSpheres | ThermoFisher Scientific | Cat#T10711 |
| NeutrAvidin | ThermoFisher Scientific | Cat#31000 |
| Protocatechuate dioxygenase | Sigma-Aldrich | Cat#P8279 |
| Protocatechuic acid | Sigma-Aldrich | Cat#03930590 |
| Propyl gallate | Sigma-Aldrich | Cat#02370 |
| Trolox | Sigma-Aldrich | Cat#238813 |
| 4-nitrobenzyl alcohol | Sigma-Aldrich | Cat#N12821 |
| Hexokinase | Sigma-Aldrich | Cat#H4502 |
| Acetyl-CoA | Sigma-Aldrich | Cat#A2056 |
| Herculase II Fusion DNA Polymerase | Agilent | Cat#600675 |
| Critical commercial assays | | |
| DNA SizeSelector-I SPRI magnetic beads | Aline Biosciences | Cat#Z-6001 |
| Deposited data | | |
| Single-molecule fluorescence source data | Zenodo | DOI: 10.5281/zenodo.16687840 |
| Experimental models: Organisms/strains | | |
| *S. cerevisiae* MATa, ura3-1, leu2-3,112, trp1-1, his3-11,15, ade2-1, pep4∆::HIS3, prb∆::his3, prc1∆::hisG | Rosen et al.^50^; Inada et al.^103^ | YF702 |
| *S. cerevisiae* MATa, ura3-1, leu2-3,112, trp1-1, his3-11,15, ade2-1, pep4∆::HIS3, prb∆::his3, prc1∆::hisG, RPB1-SNAPf::NATMX, MED7-HA3-DHFR::Hygromycin^R^ | This study | YSB3499 |
| *S. cerevisiae* MATa, ura3-1, leu2-3,112, trp1-1, his3-11,15, ade2-1, pep4∆::HIS3, prb∆::his3, prc1∆::hisG, RPB1-DHFR::Hygromycin^R^, MED7-HA3-SNAPf::KanMX | This study | YSB3613 |
| *S. cerevisiae* MATa, ura3-1, leu2-3,112, trp1-1, his3-11, 15, ade2-1, pep4∆::HIS3, prb∆::his3, prc1∆::hisG, RPB1-HA3-HALO::NATMX, MED7-HA3-SNAPf::KanMX | Baek et al.^95^ | YSB3687 |
| Oligonucleotides | | |
| fSNAP_gs_GCN4_F (O4184): gggctgggtggctctggcggttccggctcaATGTCCGAATATCAGCCAA | Integrated  DNA Technologies | N/A |
| GCN4_pRJR_R (O4185): cgggggatgcgggtccggtcgcgccccCTACAGAGAAACTTCTTCAGTGGATT | Integrated  DNA Technologies | N/A |
| fSNAP_gs_R (O4081): accgccagagccACCCAGCCC | Integrated  DNA Technologies | N/A |
| Amp-gibson_F (O4018): CGGATGGCATGACAGTAAGAGAATTATGCAGTGCTGCCAT | Integrated  DNA Technologies | N/A |
| pRJR_F (O4082): GGGGCGCGACCGGACCCGCA | Integrated  DNA Technologies | N/A |
| Amp-gibson_R (O4019): ATGGCAGCACTGCATAATTCTCTTACTGTCATGCCATCCG | Integrated  DNA Technologies | N/A |
| P830: ACCTTAAGAACCGGATCACAATCTCCTCCATCGTCGTCCCATACCCATACGATGTTCCT | Integrated  DNA Technologies | N/A |
| P831: TGAAAATGTATATAGTTACACAAATGATTTATAATAGTGTGGGCGGCGTTAGTATCGAAT | Integrated  DNA Technologies | N/A |
| upstream primer:  biotin-TTGGGTAACGCCAGGGT | Integrated  DNA Technologies | N/A |
| downstream primer:  Alexa488-AGCGGATAACAATTTCACACAG | Integrated  DNA Technologies | N/A |
| downstream primer 2:  Alexa488-CGAGATCCTCTAGAGTCGG | Integrated  DNA Technologies | N/A |
| DNA template, see Table S4 | This study | N/A |
| Recombinant DNA | | |
| pSH556 (Gal4-Gcn4) (BE473) | Reeves and Hahn^104^ | N/A |
| pRJR-Gal4-fSNAP-vp16 (BE559) | Jeon et al.^105^ | N/A |
| pRJR-Gal4-fSNAP-Gcn4 (BE573) | This study | N/A |
| pRJR-Gal4vp16 (BE165) | Cho et al.^106^ | N/A |
| pUC18-G5CYC1 G- (SB649) | Johnson et al.^107^ | N/A |
| pUC18-5xGal4UASmut-CYC1 G- (SB1958) | This study | N/A |
| pBS-SKII-3XHA-eDHFR-Hygromycin (YV317) | Baek et al.^95,108^ | N/A |
| pBS-SKII-3XHA-fSNAP-Kan (YV311) | Baek et al.^95^ | N/A |
| Software and algorithms | | |
| Glimpse | Gelles Lab | https://github.com/gelles- brandeis/Glimpse |
| LabView | National Instruments | https://www.ni.com/en-us.html |
| MATLAB | The MathWorks | https://www.mathworks.com/ |
| Imscroll | Friedman and Gelles^109^ | https://github.com/gelles-  brandeis/CoSMoS_Analysis |
| Tapqir | Ordabayev et al.^110^ | https://github.com/gelles-brandeis/tapqir |
| Matlab code to analyze single-molecule fluorescence intensity distributions | This study | Doi: 10.5281/zenodo.16687840 |
| Other | | |
| Homogenizer | Wheaton | Cat#62400-802 |
