## Supplemental figures and tables for "A mechanism of synergistic Mediator recruitment in RNA polymerase II transcription activation revealed by single-molecule fluorescence"

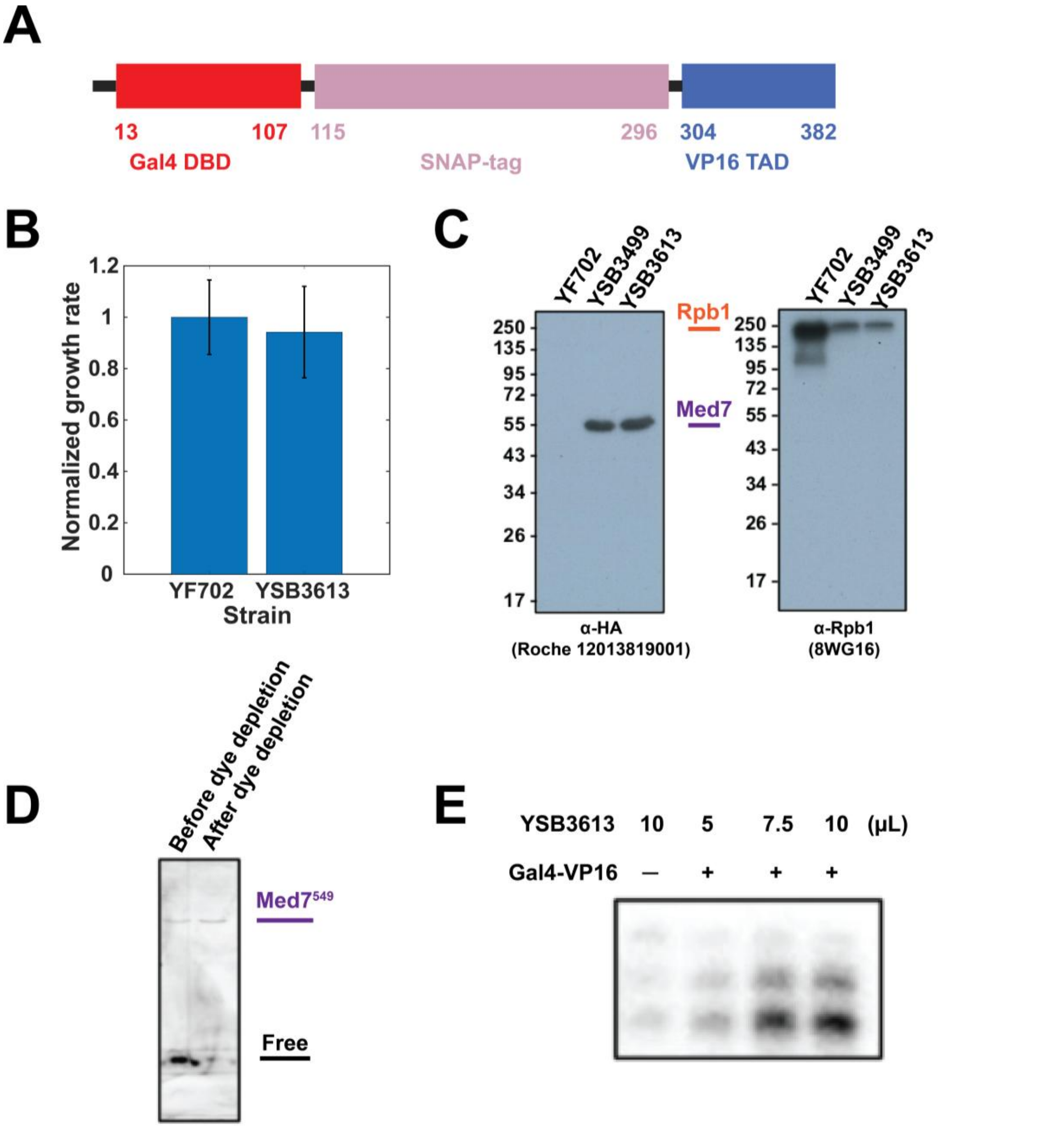


**Figure S1. TA construct and yeast nuclear extract activities, related to Figure 1. (A)** Domain structure of Gal4-SNAP-VP16 construct [S1] (numbers represent amino acid residues). **(B)** Growth rates (±S.E.M.; N = 5) of untagged (YF702) and tagged (YSB3613; Med7-HA-SNAP, Rpb1-DHFR) strains, normalized to that of the untagged strain. **(C)** Western blots of yeast nuclear extracts (YNE) prepared from the strains used in this work (**Table S1**) using antibodies against HA-epitope tag (left, for detection of Med7-HA fusions) and against Rpb1 (right). **(D)** YNE containing $Med7^{549}$ before and after removal of unincorporated SNAP-Surface-549 dye, analyzed by fluorescence scan of an SDS-PAGE gel. **(E)** Bulk in vitro transcription assay showing incorporation of α-[$^{32}$P]UTP into RNA products by YNE containing $Med7^{549}$ and Rpb1-DHFR. Transcription activity is Gal4-VP16-dependent and increases with the addition of increasing volumes of YNE.

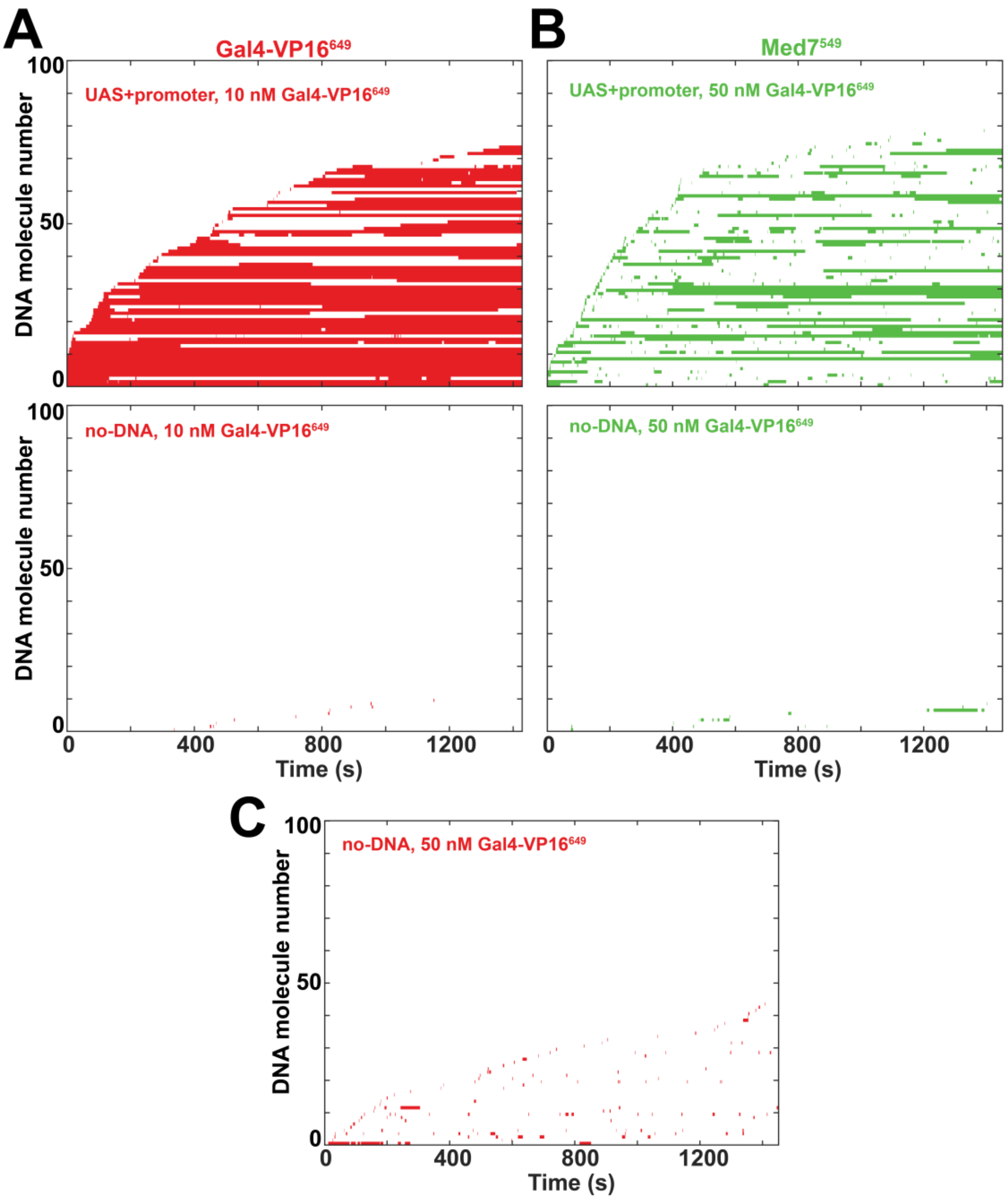


**Figure S2. Specificity of TA and Mediator binding for DNA molecules, related to Figure 2.**
**(A)** *Top:* Rastergram plot for Gal4-VP16$^{649}$ binding at UAS+promoter DNA molecules in the experiment at 10 nM Gal4-VP16$^{649}$ (**Figure 2A). **Plot shows data from 100 randomly selected DNA locations, sorted by the time of first Gal4-VP16$^{649}$ arrival. Each horizontal line displays the time record for a single DNA location, color coded to indicate bound Gal4-VP16$^{649}$ presence (color) or absence (white). *Bottom:* same as top, but for 100 randomly selected control no-DNA locations in the same experiment. **(B)** Same as (A) but for Med7$^{549}$ in the experiment at 50 nM Gal4-VP16$^{649}$ (**Figure 2C**). **(C)** Rastergram plot for Gal4-VP16$^{649}$ binding at 100 randomly selected control no-DNA locations in the experiment at 50 nM Gal4-VP16$^{649}$ (**Figure 2C).**

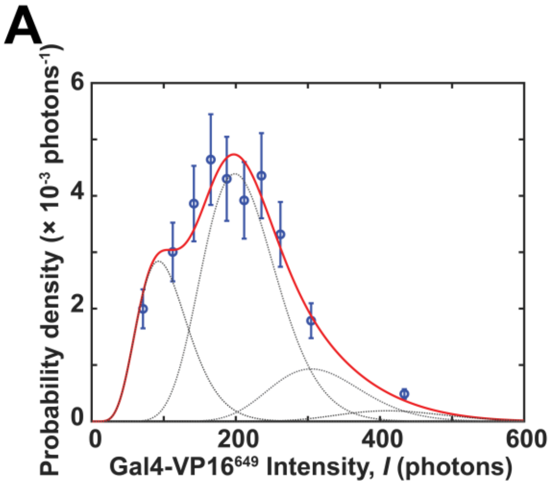


| Parameter | Value (± SE) |
|---|---|
| $\mu_1$ | 106 ± 13 (photons) |
| $w$ | 0.076 ± 0.02 (photons$^{-1}$) |
| $P_1$ | 0.25 ± 0.17 |
| $P_2$ | 0.57 ± 0.15 |
| $P_3$ | 0.15 ± 0.06 |
| $P_4$ | 0.03 ± 0.03 |

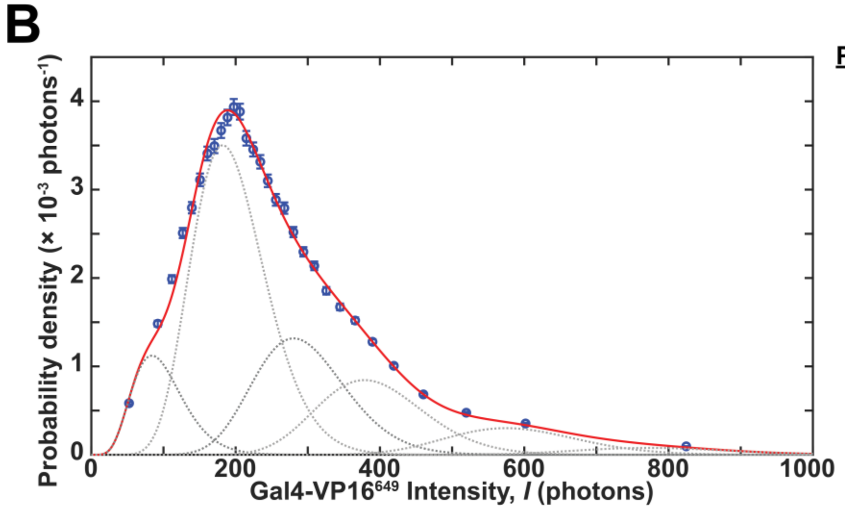


| Parameter | Value (± SE) |
|---|---|
| $\mu_1$ | 98 ± 0.5 (photons) |
| $w$ | 0.071 ± 0.002 (photons$^{-1}$) |
| $P_1$ | 0.10 ± 0.003 |
| $P_2$ | 0.45 ± 0.01 |
| $P_3$ | 0.21 ± 0.01 |
| $P_4$ | 0.16 ± 0.01 |
| $P_5$ | 0.0004 ± 0.01 |
| $P_6$ | 0.07 ± 0.01 |
| $P_7$ | 0 ± 0.005 |
| $P_8$ | 0.02 ± 0.002 |

**Figure S3. Single-DNA Gal4-VP16$^{649}$ fluorescence intensity distributions from the experiment at 10 nM Gal4-VP16$^{649}$, related to Figures 2, 5, 6 and S7. (A)** *Left*, Distribution of fluorescence intensities (blue) of the first Gal4-VP16$^{649}$ molecule to bind at each DNA location (± S.E.; $N_{DNA}$ = 506). Also shown is the fit to a gamma distribution mixture model (red) which is the sum of four components (gray) corresponding to 1, 2, 3, or 4 dye molecules arriving with the Gal4-VP16 molecule(s) (see Methods). *Right*, Parameter values (± S.E.) from the fit: $\mu_1$, mean fluorescence intensity of a single dye moiety; $w$, Gamma component inverse scale parameter; $P_1$, $P_2$, $P_3$, and $P_4$, component fractional amplitudes. **(B)** *Left*, Distribution of fluorescence intensities of equilibrium Gal4-VP16$^{649}$ dwells and fit to an eight-component gamma distribution mixture model plotted as in (A). *Right*, Parameter values from the fit. Numbers of observations, distributions and fit plots, and parameter values for all experiments in **Figures 5A, 6,** and **S7** are reported in **Supplementary Data File 1**.

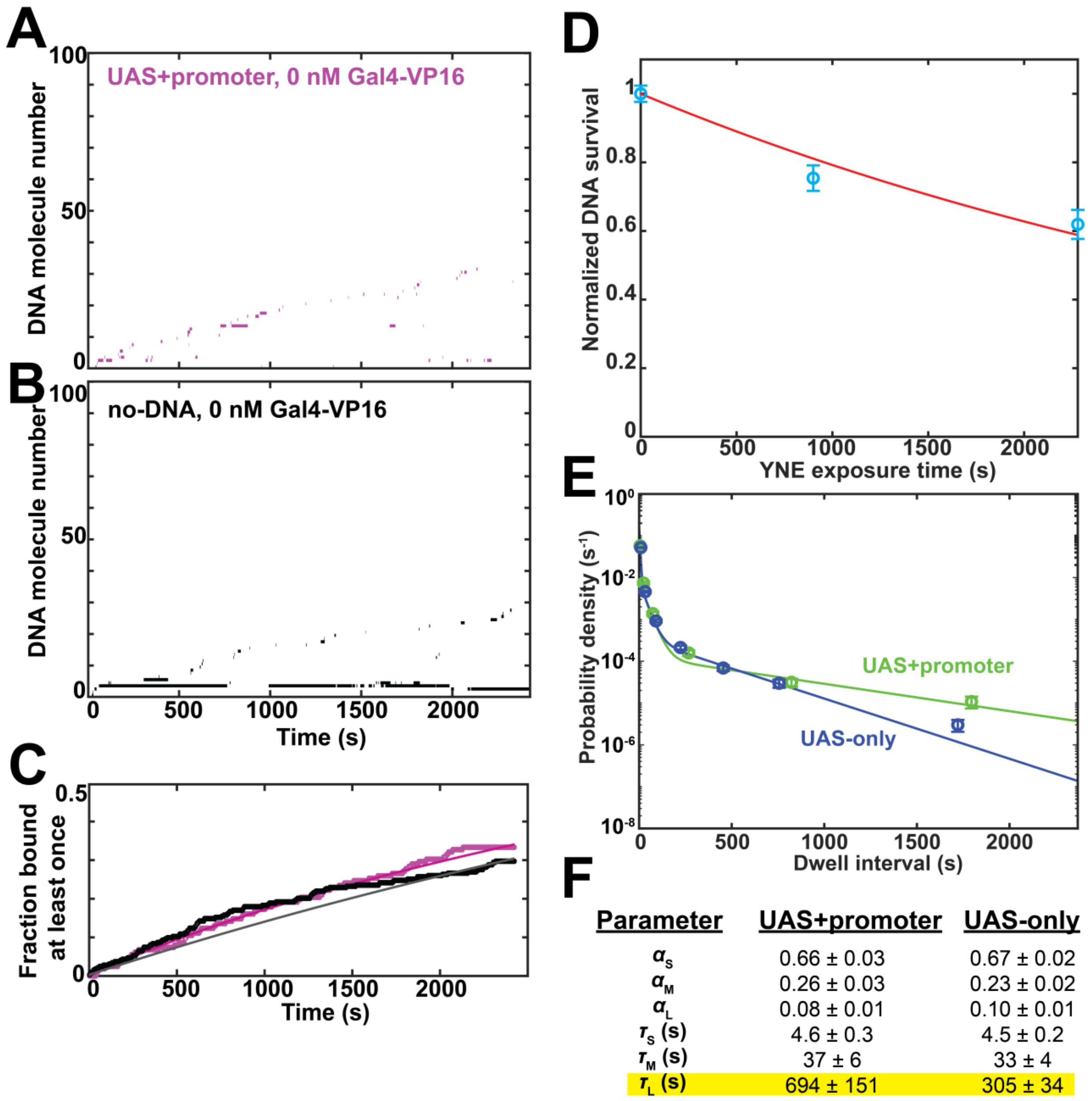


| Parameter | UAS+promoter | UAS-only |
|---|---|---|
| $\alpha_S$ | 0.66 ± 0.03 | 0.67 ± 0.02 |
| $\alpha_M$ | 0.26 ± 0.03 | 0.23 ± 0.02 |
| $\alpha_L$ | 0.08 ± 0.01 | 0.10 ± 0.01 |
| $\tau_S$ (s) | 4.6 ± 0.3 | 4.5 ± 0.2 |
| $\tau_M$ (s) | 37 ± 6 | 33 ± 4 |
| $\tau_L$ (s) | 694 ± 151 | 305 ± 34 |

**Figure S4. Further analysis of and control experiments for Mediator binding studies, related to Figure 3. (A, B, C)** Med7[549] binding to UAS+promoter DNA or control no-DNA locations in the absence of Gal4-VP16, performed as a control experiment for the experiment shown in **Figure 3A** and **Figure 3F**, green. Rastergrams (A, B) are plotted as in **Figure 3A, E**, respectively; time to first Med7[549] binding distributions (C) are plotted and fit as in **Figure 3F**. Fit parameters are reported in **Table S2**. In the absence of activator, there is no appreciable difference between the two curves in (C), indicating that there is no DNA-specific binding when recombinant activator is not added. **(D)** Surface-tethered DNA detachment/cleavage upon exposure to YNE. UAS+promoter DNA molecules attached to a slide surface (as illustrated in **Figure 1**) were counted in the same microscope field of view before and after exposure in the dark to YNE under the same conditions as the experiments in **Figures 2 and 3**. Separate experiments were carried out for YNE exposure times of 0, 900, and 2,280 s, and cell contents were replaced with fresh YNE reaction mixture (see Methods) just before the post-exposure counts (time delay < 60 s) with initial numbers of DNA molecules $N_{DNA}$ = 245, 168, and 139 respectively. The fractions of DNA molecules that survived the YNE incubations were normalized by the fraction that survived in the < 60 s exposure sample, plotted (± S.E.), and fit to an exponential decay model, yielding a DNA loss rate constant $k_L$ = 2.3 [1.1, 3.5] × $10^{-4}$ $s^{-1}$ [95% C.I.], roughly an order of magnitude smaller than the values of DNA occlusion rate constant $k_i$ (**Table S2**). Possible reasons for DNA loss include detachment from the slide surface or endonucleolytic DNA cleavage (which detaches the fluorophore from the surface-tethered DNA fragment). **(E)** Distributions of DNA-specific $Med7^{549}$ dwell times on UAS+promoter (green) and UAS-only (blue) DNAs. Data are the same as in **Figure 3J** but are plotted as non-cumulative binned probability densities (circles; ± S.E.) for intervals in which one or more $Med7^{549}$ were present. Also shown are maximum likelihood fits (lines) to a three-exponential model that accounts for the separately measured contribution of non-specific surface binding (see Methods). **(F)** Parameters (± S.E.) for the fits in (E) which consist of the characteristic amplitudes ($a_S$, $a_M$, and $a_L$) and lifetimes ($\tau_S$, $\tau_M$, and $\tau_L$) of the three components. Only the $\tau_L$ value (highlighted) clearly differs between the two DNAs.

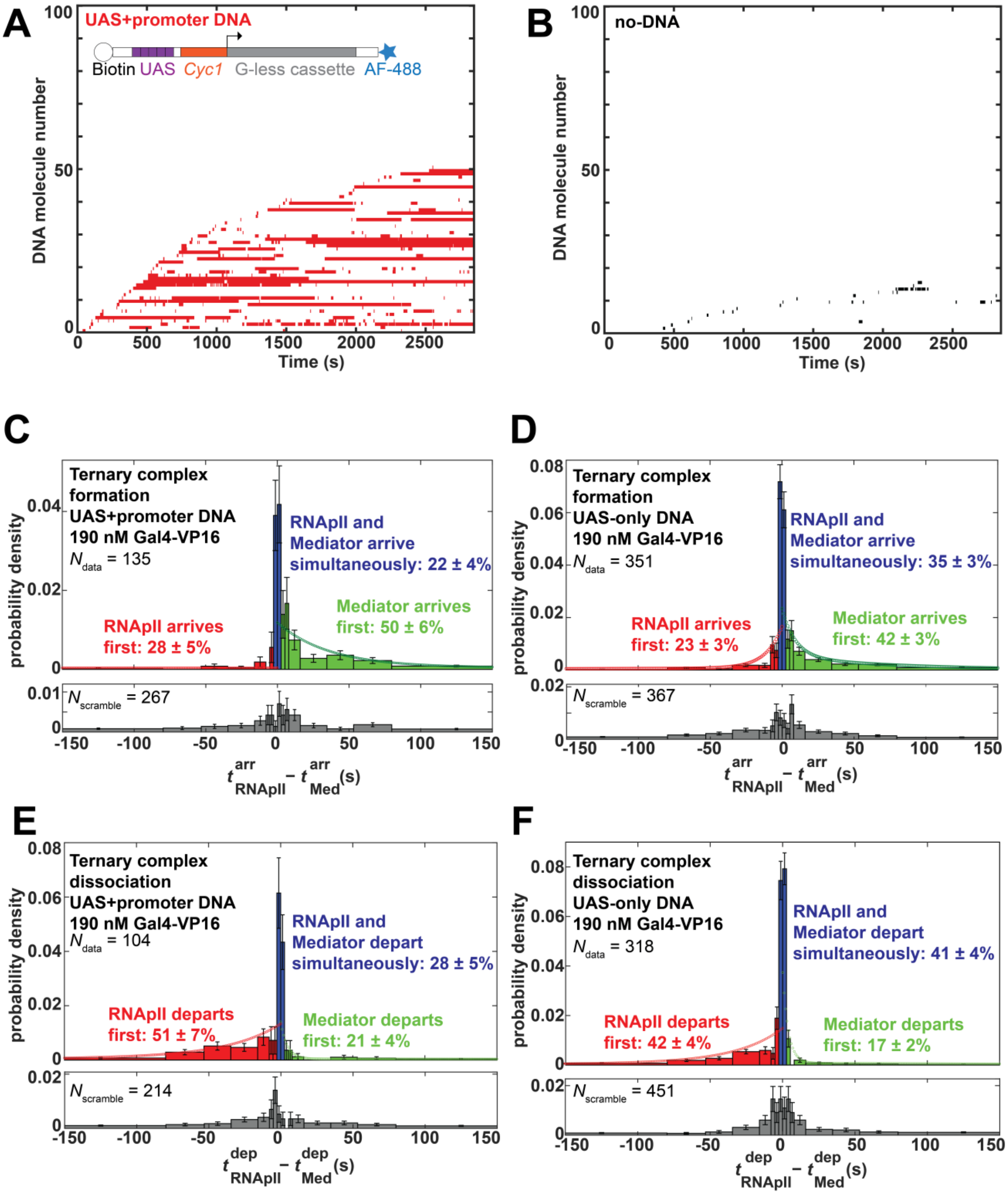


**Figure S5. Kinetic analysis of correlated DNA binding and dissociation of Med7$^{549}$ and Rpb1$^{Cy5}$, related to Figure 4. (A)** Rastergram plot of Rpb1$^{Cy5}$ dwells for 100 randomly selected UAS+promoter DNA molecules from the experiment in **Figure 4**. Plot shows data from 100 randomly selected DNA locations, sorted by the time of first Rpb1$^{Cy5}$ arrival. Colored bars indicate times during which one or more Rpb1$^{Cy5}$ molecules were bound. **(B)** Same as A, but for control no-DNA locations. **(C)** Formation of ternary complexes in which both Med7$^{549}$ and Rpb1$^{Cy5}$ were simultaneously present on UAS+promoter DNA. *Top*: Distribution of ternary complex forming events (bars; ± S.E.), divided into categories in which Med7$^{549}$

arrives > 2.7 s prior to $Rpb1^{Cy5}$ arriving (green), $Rpb1^{Cy5}$ arrives > 2.7 s prior to $Med7^{549}$ (red), or events in which $Med7^{549}$ and $Rpb1^{Cy5}$ arrive simultaneously within the 2.7 s time resolution of the experiment (blue); the relative proportion (± S.E.) of each category is indicated. To determine the contribution to the apparently simultaneous arrivals of rapid successive binding events that more closely spaced than the 2.7 s time resolution, the sequential arrivals distributions (red and green bars) were independently fit to biexponential functions and extrapolated to zero time-difference (red and green curves). The majority of the apparently simultaneous arrival events are not accounted for by these fits and are thus inferred to be true simultaneous arrivals. *Bottom*: Control analysis of paired $Med7^{549}$ and $Rpb1^{Cy5}$ records from different DNA locations lacks the prominent central peak and asymmetry of the top distribution, confirming that the simultaneous and sequential ($Med7^{549}$ followed by $Rpb1^{Cy5}$) binding observed at top are not coincidental. **(D)** Same as (C), but for UAS-only DNA. **(E)** Dissolution of ternary complexes in which both $Med7^{549}$ and $Rpb1^{Cy5}$ were simultaneously present on UAS+promoter DNA. *Top*: Distribution of ternary complex dissolution events (bars; ± S.E.), categorized into events where $Med7^{549}$ departed > 2.7 s prior to $Rpb1^{Cy5}$ departure (green), $Rpb1^{Cy5}$ departed > 2.7 s prior to $Med7^{549}$ departure (red), or events in which $Med7^{549}$ and $Rpb1^{Cy5}$ depart simultaneously within the 2.7 s time resolution of the experiment (blue); the relative proportion (± S.E.) of each category is indicated. To determine the contribution to the apparently simultaneous departures of rapid successive dissociation events that are more closely spaced than the 2.7 s time resolution, the sequential departure distributions (red and green bars) were independently fit to biexponential functions and extrapolated to zero time-difference (red and green curves). The majority of the apparently simultaneous dissolution events are not accounted for by these fits and are thus inferred to be true simultaneous departures. *Bottom*: Control analysis of paired $Med7^{549}$ and $Rpb1^{Cy5}$ records from different DNA locations confirms that the simultaneous and sequential ($Rpb1^{Cy5}$ followed by $Med7^{549}$) departures observed at top are not coincidental. **(F)** Same as (E), but for UAS-only DNA.

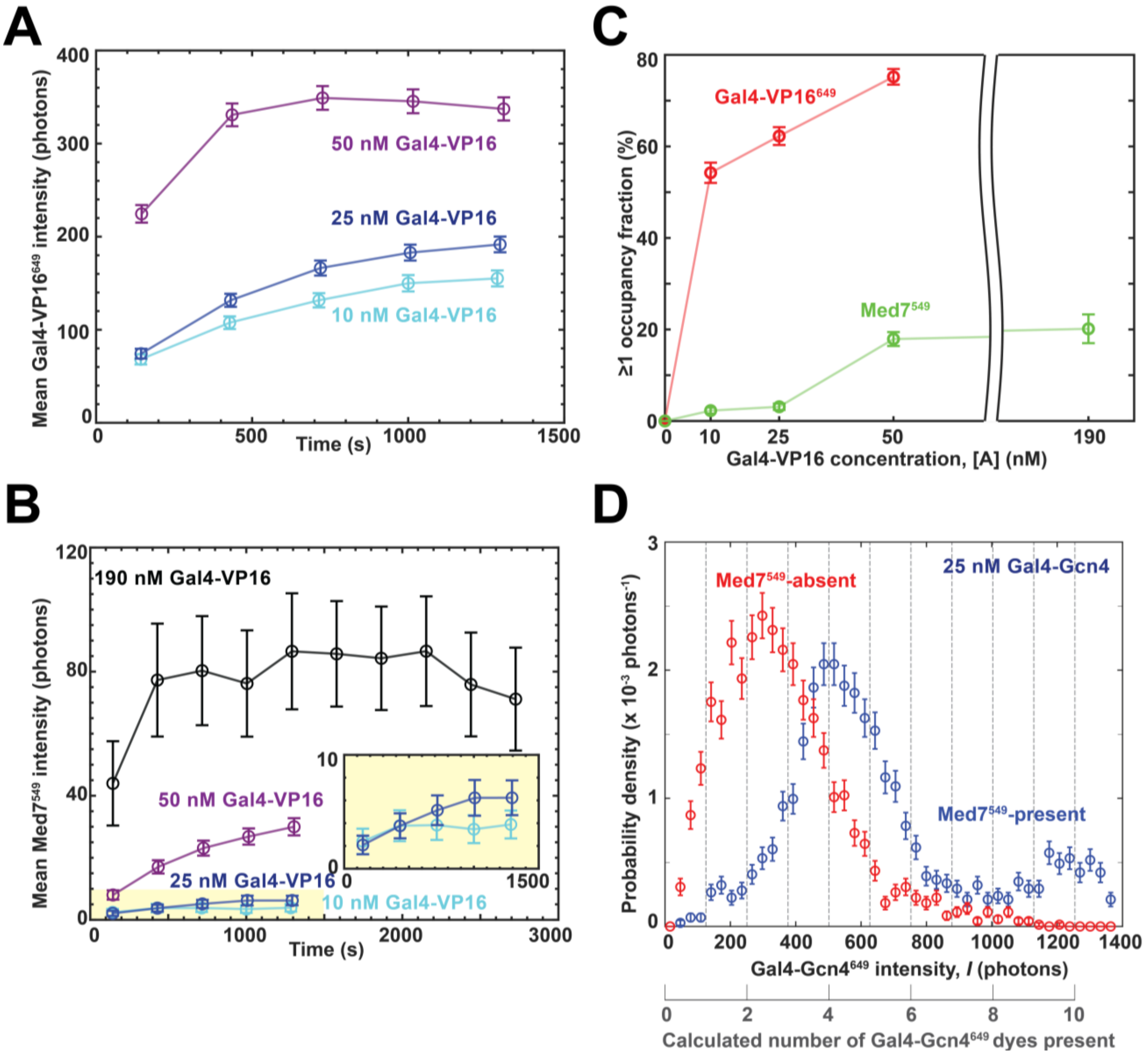


**Figure S6. Quantitation of binding of activator and Mediator to UAS+promoter DNA, related to Figure 5. (A, B)** Mean (± S.E.M.) intensities of Gal4-VP16$^{649}$ (A) and Med7$^{549}$ (B) fluorescence colocalized with DNA molecules. Each plotted point represents the time average over an interval of ~280 s centered at the point; $N_t$ and $N_{DNA}$ values are reported in **Table S3**. Inset in (B): magnified view. **(C)** Average fraction (± S.E.) of time during which a DNA molecule had at least one Gal4-VP16$^{649}$ or Med7$^{549}$ molecule, measured in a time interval (~1,100 s to ~1,400 s) at the equilibrium plateau. Where error bars are not visible, they are smaller than the points. Data in (A-C) are from the 10 and 50 nM Gal4-VP16$^{649}$ experiments in **Figure 1,** an additional analogous experiment at 25 nM, and the 190 nM unlabeled Gal4-VP16 experiment in **Figure 3A**. **(D)** Gal4-Gcn4$^{649}$ equilibrium spot intensity distributions at DNA locations in an experiment at 25 nM Gal4-Gcn4$^{649}$. Intensity distributions (± S.E.; $N_{DNA}$ = 115) are plotted for the 2,275 data points during which Med7$^{549}$ was present (blue points), or for a randomly-selected, time-matched set of 2,275 points during which Med7$^{549}$ was absent (red points). Intensities are expressed both as the number of photons detected per frame and the corresponding number of dyes present estimated from fitting intensity distributions. Data were taken from the equilibrium plateau 1,285 to 1,574 s after extract addition.

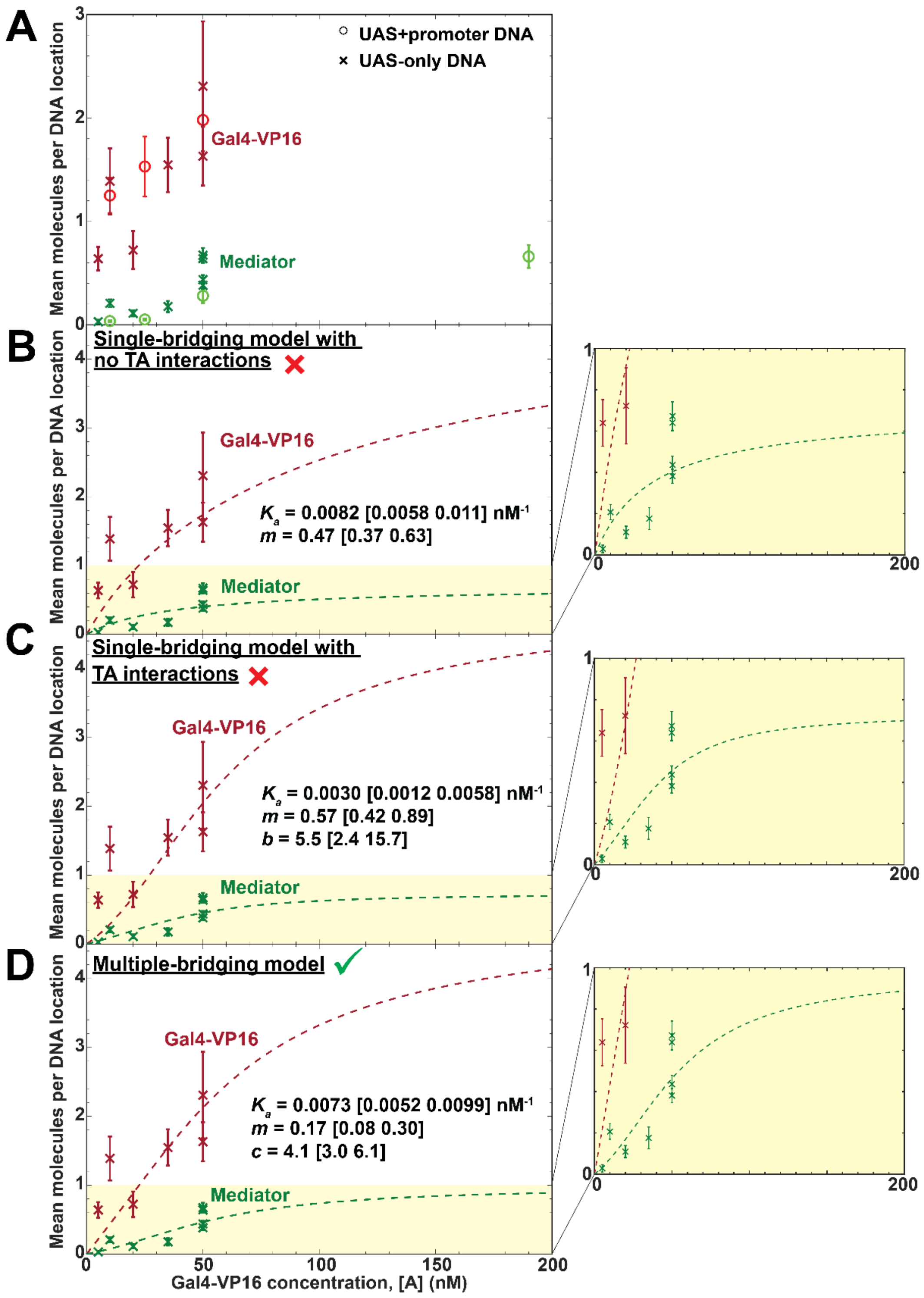


**Figure S7. Equilibrium TA and Mediator occupancy on UAS-only DNA and best fits to statistical mechanical models, related to Figure 6. (A)** Mean (± S.E.) number of Gal4-VP16 and Mediator molecules per UAS-only DNA molecule (×; each point is a separate experimental replicate; final extract protein concentration 6.6 mg /ml) compared to analogous data on UAS+promoter DNA (∘; same data as in Figure 6). Both data sets were collected in the absence of NTPs. Curve shapes with the two DNAs were quantitatively similar; the ~2-fold difference in Mediator occupancies measured is likely due to different activity levels in the two different extract preparations used in the two experiments, as seen previously[2]. Yellow shading indicates the portion of the graph shown in magnified view at right. (B,C,D) UAS-only DNA data from (A) are replotted together with the fits of combined Gal4-VP16 and Mediator data sets to the models depicted in Figures 6A, 6C, and 6E, respectively. Fits yielded the indicated parameter values [90% CIs] and respective relative likelihood values $p_{S7B} = 3.5 \times 10^{-24}$, $p_{S7C} = 7.0 \times 10^{-19}$, and $p_{S7D} = 9.9 \times 10^{-16}$. As for the UAS+promoter data (**Figure 6**), the multiple-bridging model fits the UAS-only data better than the single-bridging model with TA interactions (likelihood ratio $p_{S7D} / p_{S7C} = 1.4 \times 10^{3}$). For the UAS-only DNA, the TA and mediator data were acquired using 633 nm and 532 nm excitation (0.05 and 0.2 mW, respectively) at a frame rate of 1 frame every 2.7 s (1s exposure for each channel, plus switching times) using extract from strain YSB3687 (see **Table S1**).

| Table S1. Yeast strains and nuclear extracts, related to STAR Methods | | | | |
|---|---|---|---|---|
| Strain | Genotype | Source | Tagged proteins | Concentration (mg protein / mL)* |
| YF702 | MATa, ura3-1, leu2-3,112, trp1-1, his3-11,15, ade2-1, pep4Δ::HIS3, prbΔ::his3, prc1Δ::hisG | Rosen et al. [S2]; Inada et al. [S3] | none (wild type extract) | |
| YSB3499 | MATa, ura3-1, leu2-3,112, trp1-1, his3-11,15, ade2-1, pep4Δ::HIS3, prbΔ::his3, prc1Δ::hisG, RPB1-SNAPf::NATMX, MED7-HA3-DHFR::Hygromycin$^R$ | This study | Rpb1-SNAP Med7-HA-DHFR | 5.4 |
| YSB3613 | MATa, ura3-1, leu2-3,112, trp1-1, his3-11,15, ade2-1, pep4Δ::HIS3, prbΔ::his3, prc1Δ::hisG, RPB1-DHFR::Hygromycin$^R$, MED7-HA3-SNAPf::KanMX | This study | Med7-HA-SNAP Rpb1-DHFR | 8.8 |
| YSB3687 | MATa, ura3-1, leu2-3,112, trp1-1, his3-11, 15, ade2-1, pep4Δ::HIS3, prbΔ::his3, prc1Δ::hisG, RPB1-HA3-HALO::NATMX, MED7-HA3-SNAPf::KanMX | Baek et al. [S4] | Med7-HA-SNAP RBP1-HA-HALO | 6.6 |

*Final extract protein concentration in the single-molecule experiments.

**Table S2. Summary of Mediator binding kinetics at zero and 190 nM Gal4-VP16, related to Figure 3**[a]

| Extract | Replicate number | $k_{eff}$ (× $10^{-3}$ $s^{-1}$) | $k_i$ (× $10^{-3}$ $s^{-1}$) | $k_n$ (× $10^{-3}$ $s^{-1}$) | $k_{on}$ (× $10^{-3}$ $s^{-1}$) | $A_f$ |
|---|---|---|---|---|---|---|
| **Med7 binding to UAS+promoter DNA, −NTPs, 190 nM Gal4-VP16** | | | | | | |
| Med7-SNAP[549]/ Rpb1-DHFR | 1[b] | 1.58 ± 0.27<br>($N_{DNA}$ = 163) | 1.0 ± 0.5 | 0.09 ± 0.01<br>($N_{no\text{-}DNA}$ = 298) | 2.5 ± 0.4<br>($N_{DNA}$ = 163) | 0.63 ± 0.05 |
| | 2[d] | 2.73 ± 0.51<br>($N_{DNA}$ = 225) | 1.8 ± 0.9 | 0.17 ± 0.01<br>($N_{no\text{-}DNA}$ = 465) | 4.5 ± 0.8<br>($N_{DNA}$ = 225) | 0.61 ± 0.04 |
| | 3 | 2.91 ± 0.52<br>($N_{DNA}$ = 263) | 1.6 ± 0.9 | 0.25 ± 0.02<br>($N_{no\text{-}DNA}$ = 397) | 4.5 ± 0.8<br>($N_{DNA}$ = 263) | 0.65 ± 0.04 |
| Rpb1-SNAP[549]/ Med7-DHFR | 1 | 2.02 ± 0.44<br>($N_{DNA}$ = 267) | 1.2 ± 0.8 | 0.21 ± 0.02<br>($N_{no\text{-}DNA}$ = 249) | 3.3 ± 0.7<br>($N_{DNA}$ = 267) | 0.62 ± 0.05 |
| | 2 | 1.31 ± 0.20<br>($N_{DNA}$ = 373) | 0.9 ± 0.4 | 0.26 ± 0.02<br>($N_{no\text{-}DNA}$ = 295) | 2.2 ± 0.3<br>($N_{DNA}$ = 373) | 0.59 ± 0.04 |
| | 3 | 2.70 ± 0.39<br>($N_{DNA}$ = 510) | 2.0 ± 0.7 | 0.62 ± 0.04<br>($N_{no\text{-}DNA}$ = 360) | 4.7 ± 0.6<br>($N_{DNA}$ = 510) | 0.58 ± 0.04 |
| Weighted mean[g] | | **1.78 ± 0.10** | **1.2 ± 0.2** | | | |
| **Med7 binding to promoter-only DNA, −NTPs, 190 nM Gal4-VP16** | | | | | | |
| Med7-SNAP[549]/ Rpb1-DHFR | 1[e] | 0.54 ± 0.43<br>($N_{DNA}$ = 136) | 2.0 ± 1.9 | 0.17 ± 0.01<br>($N_{no\text{-}DNA}$ = 465) | 2.5 ± 1.8<br>($N_{DNA}$ = 136) | 0.22 ± 0.07 |
| | 2 | 0.71 ± 6<br>($N_{DNA}$ = 136) | 5.8 ± 60 | 0.25 ± 0.02<br>($N_{no\text{-}DNA}$ = 397) | 6.5 ± 60<br>($N_{DNA}$ = 136) | 0.11 ± 0.05 |
| Rpb1-SNAP[549]/ Med7-DHFR | 1 | 0.34 ± 0.36<br>($N_{DNA}$ = 196) | 1.9 ± 2 | 0.21 ± 0.02<br>($N_{no\text{-}DNA}$ = 249) | 2.2 ± 2.2<br>($N_{DNA}$ = 196) | 0.16 ± 0.06 |
| | 2 | 0.18 ± 0.87<br>($N_{DNA}$ = 372) | 2.0 ± 10 | 0.26 ± 0.02<br>($N_{no\text{-}DNA}$ = 295) | 2.2 ± 10<br>($N_{DNA}$ = 372) | 0.08 ± 0.04 |
| | 3 | 1.12 ± 0.79<br>($N_{DNA}$ = 324) | 8.0 ± 5.2 | 0.62 ± 0.04<br>($N_{no\text{-}DNA}$ = 360) | 9.2 ± 5.2<br>($N_{DNA}$ = 324) | 0.12 ± 0.05 |
| Weighted mean[g] | | **0.47 ± 0.36** | **2.3 ± 0.9** | | | |
| **Med7 binding to UAS-only DNA, −NTPs, 190 nM Gal4-VP16** | | | | | | |
| Med7-SNAP[549]/ Rpb1-DHFR | 1[c] | 2.27 ± 0.30<br>($N_{DNA}$ = 426) | 1.4 ± 0.5 | 0.09 ± 0.01<br>($N_{no\text{-}DNA}$ = 298) | 3.7 ± 0.5<br>($N_{DNA}$ = 426) | 0.61 ± 0.03 |
| **Med7 binding to UAS+promoter DNA, −NTPs, 0 nM Gal4-VP16** | | | | | | |
| Med7-SNAP[549]/ Rpb1-DHFR | 1[f] | 0.06 ± 0.3<br>($N_{DNA}$ = 162) | 1.0 ± 2 | 0.15 ± 0.02<br>($N_{no\text{-}DNA}$ = 323) | 1.1 ± 2<br>($N_{DNA}$ = 162) | 0.06 ± 0.2 |

[a]Parameter values (± S.E.) were determined by fitting time-to-first-binding distributions to a model that accounts for non-specific binding and DNA occlusion; see Methods. Parameters: $k_{eff}$, occlusion-corrected apparent first-order association rate constant; $k_i$, inactivation rate constant; $k_n$, apparent first-order rate constant of Mediator non-specific binding to no-DNA locations on slide; $N_{DNA}$ and $N_{no\text{-}DNA}$, number of DNA and no-DNA locations fit. Also included are values reparametrized from a previous model (see Methods) for comparison: $A_f$, fraction of DNA molecules capable of binding Mediator; $k_{on}$, apparent first-order rate constant of Mediator binding to active DNA molecules. Gray highlights mark values not well determined due to the minimal binding of Mediator to promoter-only DNA or when Gal4-VP16 is not present.
[b]Fit shown in **Figure 3F**, **green**; sample of data is shown in **Figure 3A** (number of time points per DNA molecule $N_t$ = 1,068).
[c]Fit shown in **Figure 3F, blue**; data are sampled in **Figure 3B** ($N_t$ = 1,068) ); $k_{eff}$ is shown in **Figure 3H**, dark blue
[d]Fit in **Figure 3G, green**; sample of data is shown in **Figure 3C** ($N_t$ = 1,189).
[e]Fit in **Figure 3G, cyan**; sample of data is shown in **Figure 3D** ($N_t$ = 1,189).
[f]Fit in **Figure S3C**; sample of data is shown in **Figure S3A,B** ($N_t$ = 906); $k_{eff}$ is shown in **Figure 3H**, magenta.
[g]Combined data from above replicates, weighted by standard error. $k_{eff}$ values are shown in **Figure 3H** green and light blue.

| Table S3: Sample sizes in mean intensity vs. time analysis, related to Figure 5[a] | | |
|---|---|---|
| **Activator concentration (nM)** | $\mathbf{N_t}$ | $\mathbf{N_{DNA}}$ |
| 10 nM | 201 | 506 |
| 25 nM | 201 | 620 |
| 50 nM | 204 | 637 |
| 190 nM | 109 | 163 |

[a]$N_{DNA}$ and $N_t$ are the number of DNA locations and number of time points per DNA location, respectively, for each individual point plotted in **Figure S6A,B**. Data from 10, 25, and 50 nM activator experiments used labeled Gal4-VP16[649], while the data from 190 nM activator use unlabeled Gal4-VP16. The **Figure S6A,B** time records were then used to identify the time interval used to calculate the equilibrium plateau quantities reported in **Figure 5A** and **Figure S6C**.

| Table S4. DNA templates, related to Figures 1 and 3 | |
|---|---|
| **DNA template** | **Sequence** |
| **UAS+promoter**<br>**[S2, S5]**<br>**(635 bp)** | TTGGGTAACGCCAGGGTTTTCCCAGTCACGACGTTGTAAAACGACG<br>GCCAGTGCCAAGCTTGCATGCCTGCAGGTCCTCGGAGGACAGTACT<br>CCGCTCGGAGGACAGTACTCCGCTCGGAGGACAGTACTCCGCTCGG<br>AGGACAGTACTCCGCTCGGAGGACAGTACTCCGACTCTAGAGGATC<br>TCGAGGCATGTGCTCTGTATGTATATAAAACTCTTGTTTTCTTCTTTT<br>CTCTAAATATTCTTTCCTTATACATTAGGTCCTTTGTAGCATAAATT<br>ACTATACTTCTATACCTCCATACCCTTCCTCCATCTATACCACCCTA<br>CTCTCCTTTCCTCATTATTCCTCCTATTATCTTCTCCTCTTCTCTCCTT<br>CTTCTATATTTCCCAAATCTATCATCATTCACTCTCATCCCCTCTTCC<br>TTCACTCCCATTCTATTCTACTCCTTTCCCTTTCCATATCCCCTCCAC<br>CCCCCTTCCTCCCCTCTTTCAATCTTATCCCCAATCATAAAATTATCT<br>CAATTATATTCTCCTTCCATACCCCCTATCATCCTCATCCCTATCACC<br>CCCCGGGTACCGAGCTCGAATTCGTAATCATGGTCATAGCTGTTTCC<br>TGTGTGAAATTGTTATCCGCT |
| **UAS-only [S5]**<br>**(187 bp)** | TTGGGTAACGCCAGGGTTTTCCCAGTCACGACGTTGTAAAACGACG<br>GCCAGTGCCAAGCTTGCATGCCTGCAGGTCCTCGGAGGACAGTACT<br>CCGCTCGGAGGACAGTACTCCGCTCGGAGGACAGTACTCCGCTCGG<br>AGGACAGTACTCCGCTCGGAGGACAGTACTCCGACTCTAGAGGATC<br>TCG |
| **Promoter-only**<br>**(635 bp)** | TTGGGTAACGCCAGGGTTTTCCCAGTCACGACGTTGTAAAACGACG<br>GCCAGTGCCAAGCTTGCATGCCTGCAGGTC~~CTTAAAGGACAGTACT~~<br>~~TTACTTAAAGGACAGTACTTTACTTAAAGGACAGTACTTTACTTAA~~<br>~~AGGACAGTACTTTACTTAAAGGACAGTACTTTA~~ACTCTAGAGGATC<br>TCGAGGCATGTGCTCTGTATGTATATAAAACTCTTGTTTTCTTCTTTT<br>CTCTAAATATTCTTTCCTTATACATTAGGTCCTTTGTAGCATAAATT<br>ACTATACTTCTATACCTCCATACCCTTCCTCCATCTATACCACCCTA<br>CTCTCCTTTCCTCATTATTCCTCCTATTATCTTCTCCTCTTCTCTCCTT<br>CTTCTATATTTCCCAAATCTATCATCATTCACTCTCATCCCCTCTTCC<br>TTCACTCCCATTCTATTCTACTCCTTTCCCTTTCCATATCCCCTCCAC<br>CCCCCTTCCTCCCCTCTTTCAATCTTATCCCCAATCATAAAATTATCT<br>CAATTATATTCTCCTTCCATACCCCCTATCATCCTCATCCCTATCACC<br>CCCCGGGTACCGAGCTCGAATTCGTAATCATGGTCATAGCTGTTTCC<br>TGTGTGAAATTGTTATCCGCT |

The UAS containing five Gal4 binding sites is denoted by underlined text.
The *CYC1* core promoter is denoted by italicized text.
The UAS with mutated Gal4 binding sites is denoted by strikethrough text.

**Supplemental References**


S1. Jeon, J., Friedman, L.J., Zhou, D.H., Seo, H.D., Adeleke, O.A., Graham, B., Patteson, E.F., Gelles, J., and Buratowski, S. (2025). Single-molecule analysis of transcription activation: dynamics of SAGA coactivator recruitment. Nat. Struct. Mol. Biol. *32*, 675–686. https://doi.org/10.1038/s41594-024-01451-y.

S2. Rosen, G.A., Baek, I., Friedman, L.J., Joo, Y.J., Buratowski, S., and Gelles, J. (2020). Dynamics of RNA polymerase II and elongation factor Spt4/5 recruitment during activator-dependent transcription. Proc. Natl. Acad. Sci. *117*, 32348–32357. https://doi.org/10.1073/pnas.2011224117.

S3. Inada, T., Winstall, E., Tarun, S.Z., Yates, J.R., Schieltz, D., and Sachs, A.B. (2002). One-step affinity purification of the yeast ribosome and its associated proteins and mRNAs. RNA *8*, 948–958. https://doi.org/10.1017/s1355838202026018

S4. Baek, I., Le, S.N., Jeon, J., Chun, Y., Reed, C., and Buratowski, S. (2022). A set of Saccharomyces cerevisiae integration vectors for fluorescent dye labeling of proteins. G3 Bethesda *12*, jkac201. https://doi.org/10.1093/g3journal/jkac201.

S5. Baek, I., Friedman, L.J., Gelles, J., and Buratowski, S. (2021). Single-molecule studies reveal branched pathways for activator-dependent assembly of RNA polymerase II pre-initiation complexes. Mol. Cell *81*, 3576-3588.e6. https://doi.org/10.1016/j.molcel.2021.07.025.
